# Cortical gene expression architecture links healthy neurodevelopment to the imaging, transcriptomics, and genetics of autism and schizophrenia

**DOI:** 10.1101/2022.10.05.510582

**Authors:** Richard Dear, Konrad Wagstyl, Jakob Seidlitz, Ross D. Markello, Aurina Arnatkevičiūtė, Kevin M. Anderson, Richard A.I. Bethlehem, Lifespan Brain Chart Consortium, Armin Raznahan, Edward T. Bullmore, Petra E. Vértes

**Affiliations:** Department of Psychiatry, University of Cambridge, Cambridge, UK; Lifespan Brain Institute, The Children’s Hospital of Philadelphia and Penn Medicine, Philadelphia, PA, USA; Department of Child and Adolescent Psychiatry and Behavioral Science, The Children’s Hospital of Philadelphia, Philadelphia, PA, USA; Department of Psychiatry, University of Pennsylvania, Philadelphia, PA, USA; McConnell Brain Imaging Centre, Montreal Neurological Institute, McGill University, Montreal, Quebec, Canada; Turner Institute for Brain and Mental Health, Monash University, Melbourne, Australia; Department of Psychology, Yale University, New Haven, CT, USA; Wellcome Centre for Human Neuroimaging, London, UK; Section On Developmental Neurogenomics, National Institute of Mental Health, Bethesda, MD, USA

## Abstract

Human brain organisation involves the coordinated expression of thousands of genes. For example, the first principal component (C1) of cortical transcription identifies a hierarchy from sensorimotor to association regions. Here, optimised processing of the Allen Human Brain Atlas revealed two new components of cortical gene expression architecture, C2 and C3, which are distinctively enriched for neuronal, metabolic and immune processes, specific cell-types and cytoarchitecture, and genetic variants associated with intelligence. Using additional datasets (PsychENCODE, Allen Cell Atlas, and BrainSpan), we found that C1-C3 represent generalisable transcriptional programmes that are coordinated within cells, and differentially phased during foetal and postnatal development. Autism spectrum disorder and schizophrenia were specifically associated with C1/C2 and C3, respectively, across neuroimaging, differential expression, and genome-wide association studies. Evidence converged especially in support of C3 as a normative transcriptional programme for adolescent brain development, which can lead to atypical supra-granular cortical connectivity in people at high genetic risk for schizophrenia.

## Introduction

How does the complex anatomical and functional organisation of the human brain develop from the expression of over twenty thousand genes ^1^? And how does this process go awry in neurodevelopmental disorders? In the last 10 years, whole-brain, whole-genome transcriptional atlases, such as the Allen Human Brain Atlas (AHBA) ^2^, have suggested that healthy brain organisation may depend on “transcriptional programmes” representing the coordinated expression of large numbers of genes over development ^3–7^.

In 2012, Hawrylycz *et al*. showed that principal components of the AHBA dataset capture distinct features of cortical anatomy ^2^. In 2018, Burt *et al*. argued that the first principal component of cortical gene expression (C1) reflects an anterior-to-posterior “neuronal hierarchy”, defined in macaque tract-tracing data by feedforward and feedback axonal connections between cortical areas ^8–10^ and indexed in humans by the ratio of T1- and T2-weighted MRI signals, a putative marker of cortical myelination ^8^. These discoveries echoed prior findings from studies of embryonic development of chick, mouse and human brains where spatially patterned transcriptional gradients have been shown to organise neurodevelopmental processes such as areal differentiation, axonal projection, and cortical lamination ^6,11–13^. Single-cell RNA sequencing data has also revealed an anterior-to-posterior gradient in the gene expression of inhibitory interneurons, which is conserved across multiple species including humans ^14^. It is therefore likely that the principal component of gene expression in the adult human cortex represents a transcriptional programme key to its normative development.

However, it is not clear that C1 is the only component of spatially patterned and neurodevelopmentally coordinated gene expression in the human brain. Hawrylycz *et al.* suggested that principal component analysis (PCA) of a restricted set of 1,000 genes in one of the six brains of the AHBA dataset revealed multiple biologically-relevant components ^2^ (**Supplementary Fig. S1**). Later, Goyal *et al.* used nonlinear dimension reduction across whole-genome spatial expression, again from only one of the six AHBA brains, to show that aerobic glycolysis was associated with a second transcriptional component ^15^. To our knowledge, more recent studies using all available AHBA data have reliably found only C1 ^8,16^. This first component has been linked to a general “sensorimotor-association axis” of brain organisation ^10^ derived from several macro-scale brain phenotypes, including among others the principal gradient of functional connectivity ^17^, maps of brain metabolism and blood flow ^15^, and the map of human cortical expansion compared to other primates ^18^. Although it is parsimonious to assume that such diverse brain phenotypes could all be determined by a single transcriptional programme, it seems more realistic to expect that multiple transcriptional programmes are important for human brain development, as is generally the case for brain development in other species ^19^.

Here we present two higher-order components of human cortical gene expression, C2 and C3, that likely represent additional transcriptional programmes distinct from the C1 component already reliably described ^8^. These higher-order components only emerged when optimised data-filtering and dimension-reduction methods were applied to the AHBA dataset. We found that C2 and C3 are each specifically enriched for biologically-relevant gene sets, and spatially co-located with distinct clusters of neuroimaging phenotypes or macro-scale brain maps. Leveraging independent RNA sequencing datasets on single-cell and developmental gene expression, we further demonstrated that all three components are generalisable to other datasets, representative of coordinated transcription within cells of the same class, and dynamically differentiated over the course of foetal, childhood and adolescent brain development. Finally, by triangulating evidence across case-control neuroimaging, differential gene expression, and genome-wide association studies (GWAS), we demonstrated that components C1 and C2 are specifically associated with autism spectrum disorder (ASD), and C3 with schizophrenia. While prior studies have used the AHBA to derive gene sets correlated with disorder-related MRI phenotypes ^20–25^, this disorder-first, “imaging transcriptomics” ^26–28^ approach is susceptible to identifying genes whose co-location with MRI phenotypes reflects secondary associations or consequences of a disorder, such as behavioural changes (e.g. smoking, alcohol use), physical health disorders (e.g. obesity, diabetes), or pharmacological treatment ^29–31^. What is of most interest for neurodevelopmental disorders is to understand the pathogenic provenance of a clinically diagnosable disorder – to ask “what developed differently?” rather than merely “what is different?”. Our approach sought to distinctively address the question of what “develops differently” based on an understanding of “normal development”, by linking genetic risks and atypical phenotypes to a generalisable transcriptional architecture of healthy brain development.

## Results

### Three components pattern cortical gene expression

We first applied PCA to the entire AHBA dataset of 6 adult brains ^2^. Microarray measurements of relative messenger RNA levels were processed to represent mean expression of ∼16,000 genes at each of the 180 regions of the left hemispheric cortex defined by the HCP-MMP parcellation ^32–34^ (**Methods**). We initially found that higher-order components (C2, C3) estimated by PCA of the resulting {180 × 16,000} data matrix were not robust to sampling variation of the six donor brains, with low generalisability, *g*, compared to C1: *g*_C1_ = 0.78*, g*_C2_ = 0.09*, g*_C3_ = 0.14 (**Methods**). However, two data processing improvements were found to enhance the generalisability of higher order components. First, we optimised the trade-off involved in excluding noisy data – by filtering spatially inconsistent genes (with low differential stability ^35^) and under-sampled brain regions – while seeking to maximise the anatomic and genomic scope of the data matrix (**Extended Data Fig. 1**). Second, we used the non-linear dimension reduction technique of diffusion map embedding (DME), instead of linear PCA, to identify coordinated gene expression patterns from the matrix. DME is robust to noise and more biologically plausible than PCA in this context because of its less strict orthogonality constraints (**Methods**). We found that while PCA and DME both identified the same components from the filtered gene expression matrix (**Extended Data Fig. 1d**), using DME was necessary to achieve high generalisability *g* while also retaining sufficient genes for downstream enrichment analyses.

We applied DME to the (137 × 7,937} filtered AHBA data matrix comprising the expression of the 50% most stable genes measured in the 137 cortical areas with data available from at least three brains. The generalisability of the first three components was substantially increased, i.e., *g*_C1_ = 0.97, *g*_C2_ = 0.72, *g*_C3_ = 0.65, while the generalisability of even higher-order components remained low, e.g., *g*_C4_ = 0.28 (**Fig 1a**). We found that the cortical maps of C2 and C3 derived from DME on filtered data were more spatially smooth than the corresponding PCA-derived maps on unfiltered data (**Fig. 1b**), consistent with higher generalisability indicating less contamination by spatially random noise. C1-C3 were also robust to variations in parameters for processing the AHBA, including choice of parcellation template (**Extended Data Fig. 2**). Finally, the transcriptional patterns represented by C1-C3 in the AHBA dataset were reproducible in an independent PsychENCODE dataset comprising bulk RNA-seq measurements of gene expression at 11 cortical regions from N=54 healthy controls ^36^ (regional correlation *r*_C1_ = 0.85, *r*_C2_ = 0.75, *r*_C3_ = 0.73; see **Extended Data Fig. 3** and **Supplementary Table S5**).

**Figure 1:**
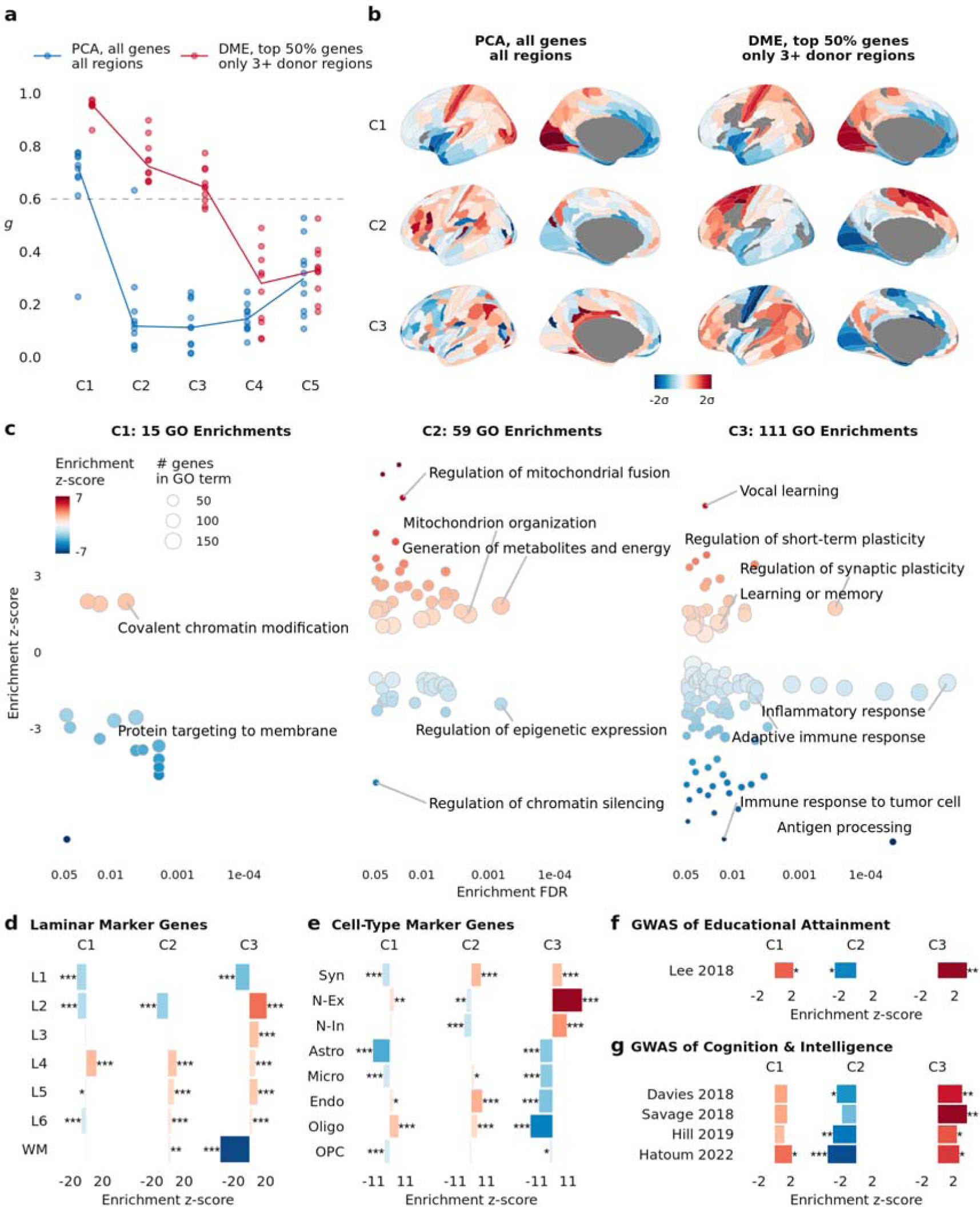
Three generalisable components of human cortical gene expression were enriched for biological processes, cytoarchitecture, and cognitive capacity. **a**, To identify robust components of cortical gene expression, we split the six-brain AHBA dataset into two disjoint triplets of three brains, applied PCA to each triplet, and correlated the resulting matched components (C1, C2, C3…) (Methods). For each component, the median absolute correlation over all 10 permutations of triplet pairs was a proxy for its generalisability, g. Using PCA and previously published best practices for processing the AHBA dataset ^33,34^, generalisability decreased markedly beyond the first component: g _C1_ = 0.78, g_C2_ = 0.09, g_C3_ = 0.14 (Fig. 1a). Using diffusion map embedding (DME) with the top 50% most stable genes, and the 137 regions with data available from at least three brains, the generalisability of the first three components substantially increased: g_C1_ = 0.97, g_C2_ = 0.72, g_C3_ = 0.65 (Fig 1a). **b**, Cortical maps of brain regional scores of components C1-C3 estimated by DME on the filtered AHBA dataset displayed smooth spatial gradients (right; Moran’s I ^37^ 0.48, 0.58, and 0.21 for C1-C3 respectively), unlike those of PCA on the unfiltered data (left; Moran’s I 0.50, 0.09 and 0.07). **c**, Gene Ontology Biological Process enrichments for C1-C3 showed that the number of significant enrichments was greater for higher-order components, illustrating that they were more biologically specific. C2-positive genes were enriched for metabolism while C2-negative genes were enriched for regulatory processes; C3-positive genes were enriched for synaptic plasticity and learning while C3-negative genes were enriched for immune processes. **d,** C1-C3 were distinctively enriched for marker genes of six cortical layers and white matter (WM) ^38^. **e**, C1-C3 were also distinctively enriched for marker genes of cell types and synapses ^44^. **f**, All three components were significantly enriched for genes mapped to common variants associated with educational attainment in prior GWAS data ^39^. **g**, C2 and C3 (but not C1) were significantly enriched for genes mapped to common variation in intelligence and cognition across four independent GWAS studies ^40–43^. For d-g, significance was computed by two-sided permutation tests (Methods) and FDR-corrected across all tests in each panel; *, **, *** respectively indicate FDR-corrected two-sided p-values: 0.05, 0.01, 0.001.

The first three DME components, C1-C3, explained 38%, 10%, and 6.5%, respectively, of the total variance of the filtered AHBA dataset (**Methods**). The proportion of variance explained was related to the number of genes that were strongly weighted (absolute correlation |*r*| ≥ 0.5) on each component: 4,867 genes (61%) were strongly weighted on C1, 967 genes (12%) on C2, and 437 genes (5.5%) on C3 (**Supplementary Fig. S2**). The three components also had distinct axial alignments in anatomical space, and the co-expression network of cortical regions displayed clear anatomical structure even when the highest-variance C1 component was regressed out (**Extended Data Fig. 4**). These findings demonstrate that these three expression patterns shared across hundreds to thousands of genes are likely to be biologically relevant.

To interpret the DME-derived components in more detail, we first used enrichment analyses of the weights of the 7,973 genes on each component (**Methods**). Many more Gene Ontology (GO) Biological Process terms were significantly enriched (with false discovery rate [FDR] = 5%) for C2 (59 GO terms) and C3 (111 GO terms) than for C1 (15 GO terms) (**Fig. 1c**).

Although C1 was enriched for relatively few, functionally general biological processes, it precisely matched the first principal component previously reported (*r* = 0.96) ^8^. The same interneuron marker genes (*SST*, *PVALB*, *VIP*, *CCK*) and glutamatergic neuronal genes (*GRIN* and *GABRA*) were strongly weighted with opposite signs (positive or negative) on C1 (**Supplementary Fig. S3**).

For genes positively-weighted on C2, 23 of 36 enrichments were for metabolic processes, and for negatively-weighted genes, 19 of 23 enrichments were for epigenetic processes (**Fig. 1c, Supplementary Table 2**). Whereas, for genes positively-weighted on C3, 19 of 27 enrichments were related to synaptic plasticity or learning, and for negatively-weighted genes, 33 of 84 enrichments involved the immune system. We further analysed enrichment for genes identified as markers of specific cortical layers ^38^ (**Fig. 1e**) and cell types ^44^ (**Fig. 1f**), and in each case observed distinct enrichment profiles for C1-C3. For example, genes positively-weighted on C3 were enriched for marker genes of neurons, synapses, and cortical layers 2 and 3 (L2, L3), whereas genes negatively-weighted on C3 were enriched for glial (especially oligodendroglial) marker genes.

We also explored the biological relevance of the three components by enrichment tests for genes associated with variation in adult cognitive capacity. We found that all three components C1-C3 were enriched for genes significantly associated with educational attainment (**Fig. 1f**) ^39^. Across four independent GWAS studies of intelligence and cognition ^40–43^, genes strongly weighted on C1 were not significantly enriched, but genes negatively-weighted on C2 were enriched for genetic variants associated with intelligence in three of the four studies, and genes positively-weighted on C3 were enriched for genes identified by all four prior GWAS studies of intelligence (**Fig. 1g**).

### Neuroimaging maps align to three transcriptional components

Prior work has linked gene transcription to a multimodal “sensorimotor-association axis” (S-A axis) ^10^ of brain organisation, defined as the composite of 10 brain maps, comprising the first principal component of gene expression (C1) and 9 other MRI or PET neuroimaging maps that were selected to differentiate sensorimotor and association cortices. We first aimed to build on this work by analysing the correlation matrix of the same set of 9 brain maps together with the three transcriptional components derived from DME of the filtered AHBA dataset. Data-driven cluster analysis of this {12 x 12} correlation matrix identified three clusters, each including one of the orthogonal transcriptional components (**Fig. 2a, Methods**). C1 was clustered together with 2 MRI maps: the T1w/T2w myelination marker ^45^ and cortical thickness ^46^; C2 was clustered with 5 maps: aerobic glycolysis ^47^, cerebral blood flow ^48^, cortical expansion in humans relative to non-human primates ^18^, inter-areal allometric scaling ^49^ and external pyramidal cell density ^50^; and C3 was clustered with 2 maps: the principal gradient of fMRI connectivity ^17^ and first principal component of cognitive terms meta-analysed by Neurosynth ^51^. While some maps were specifically aligned to one component, e.g. aerobic glycolysis r_C2_ = 0.66 (p_spin_ = 0.004, FDR < 5%), others were moderately correlated with multiple transcriptional components, e.g., for cerebral blood flow: *r*_C1_ *=* 0.25, *r*_C2_ = 0.28 and *r*_C3_ = 0.33. This clustering analysis suggests that it is overly parsimonious to align all 9 neuroimaging phenotypes with just one transcriptional component (C1) as part of a singular sensorimotor-association cortical axis.

**Figure 2:**
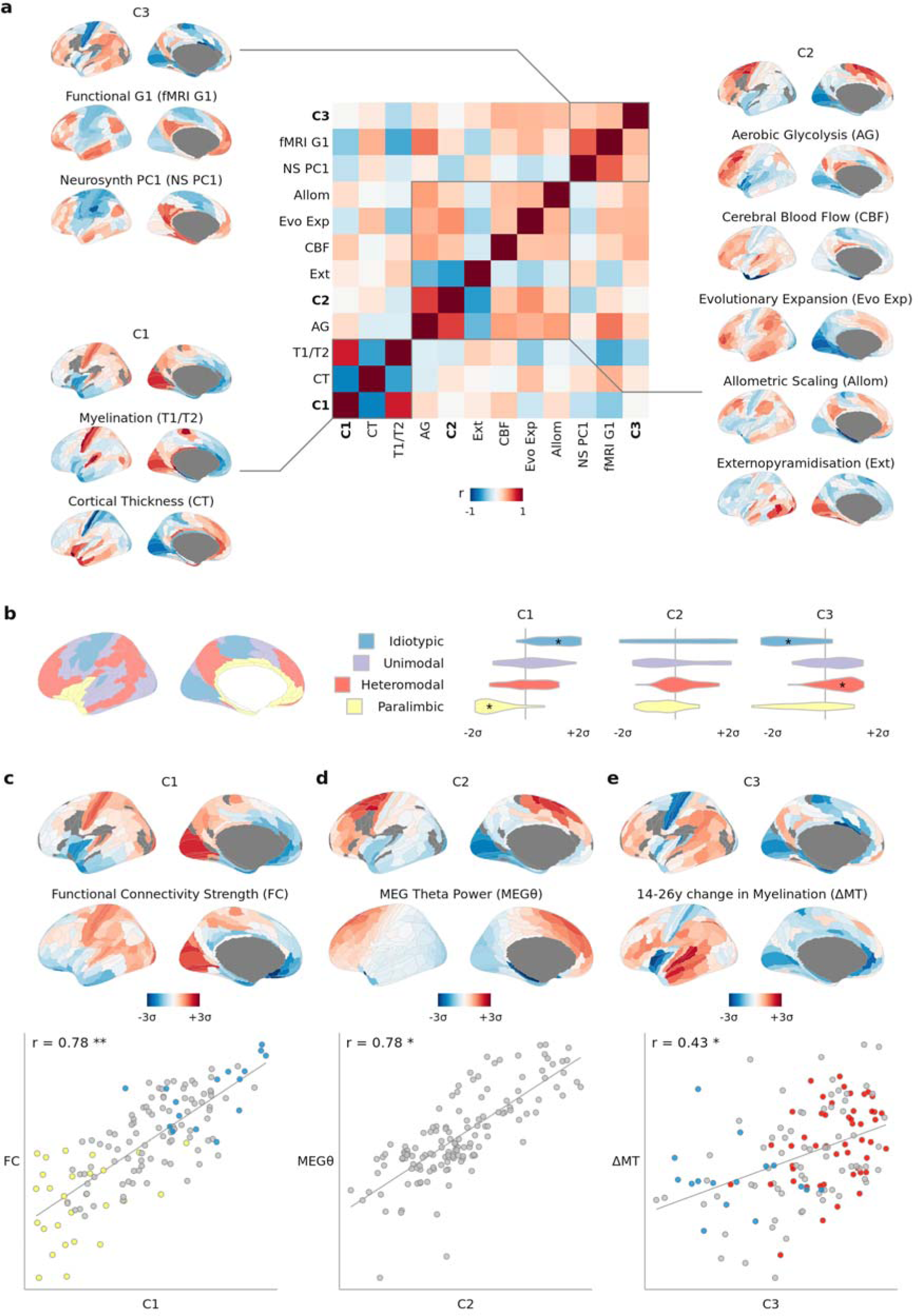
Neuroimaging and macro-scale maps of brain structure, function, and development were distinctively co-located with three components of cortical gene expression. **a**, Correlation matrix of intrinsic transcriptional components C1-C3 together with the nine neuroimaging-and physiologically-derived maps that Sydnor et al. combined with C1 to define a ‘sensorimotor-association axis’ of brain organisation ^10^. Many of the maps were not highly correlated to each other (median |r|=0.31), and data-driven clustering of the matrix revealed three distinct clusters around each of the mutually orthogonal transcriptional components C1-C3, demonstrating that all three components are relevant for understanding macroscale brain organisation. **b**, Distributions of regional scores of C1-C3 in histologically-defined regions of laminar cytoarchitecture ^52^. C1 distinguished idiotypic (p = 0.005) and paralimbic regions (p = 0.002), while C3 distinguished idiotypic (p = 0.002) and heteromodal regions (p = 0.01). * indicates FDR-adjusted two-sided p-value < 0.05, where p-value was computed by permutation test as the percentile of the mean z-score relative to null spin permutations, with adjustment for multiple comparisons across all 12 tests. **c**, Degree of fMRI functional connectivity ^53,54^ was significantly aligned to C1 (r = 0.78, p _spin_ < 0.001). Blue/yellow highlighted points correspond to idiotypic/paralimbic cytoarchitectural regions as in b. **d**, MEG-derived theta power ^55^ was significantly aligned to C2 (r = 0.78, p_spin_ = 0.002). **e**, Regional change in myelination over adolescence ^56,57^ was significantly aligned to C3 (r = 0.43, p_spin_ = 0.009). Blue/red highlighted points correspond to idiotypic/heteromodal cytoarchitectural regions as in b. For panels c-d, *, **, *** respectively indicate FDR-corrected two-sided spin-permutation p-values: 0.05, 0.01, 0.001, with corrections for multiple comparisons of all maps in panels c-d being compared with all of C1-C3.

We also found that the three transcriptional components were associated with a wider range of cellular, functional and developmental phenotypes than the 9 neuroimaging maps above, and that these associations were again distinct for the three components. For example, at cellular scale, histologically-defined regions of laminar cytoarchitectural differentiation ^52^ were co-located with C1 and C3, but not C2 (ANOVA*, p* < 0.001; **Fig. 2b**). In functional MRI and magnetoencephalography (MEG) data, we found that weighted nodal degree of cortical regions in an fMRI network ^53,54^ was strongly correlated with C1 (*r*_C1_ = 0.78, *p*_spin_ < 0.001, FDR = 5%, **Fig. 2c**) but not C2 or C3 (*r*_C2_ = −0.01, *r*_C3_ = 0.00); across all canonical frequency intervals of MEG data ^55^, an FDR-significant association was observed between theta band (4-7 Hz) oscillations and C2 (*r*_C2_ = 0.78, *p*_spin_=0.002, FDR = 5%, **Fig. 2d**) but not C1 or C3 (*r*_C2_ = −0.18, *r*_C3_ = −0.02); see **Extended Data Fig. 5** for other MEG results. And in support of the hypothetical prediction that adult brain transcriptional programmes are neurodevelopmentally relevant, we found that a prior map of adolescent cortical myelination, as measured by change in magnetisation transfer between 14-24 years (ΔMT) ^56,57^, was significantly co-located with C3 (*r*_C3_ = 0.43, *p*_spin_ = 0.009, **Fig. 2e**) but not C1 or C2 (*r*_C2_ = 0.17, *r*_C3_ = 0.15).

### C1-C3 are distinctly developing intra-cellular programmes

We next used two additional RNA-seq datasets to investigate the consistency of AHBA-derived components with gene co-expression in single cells, e.g., neurons or glia, and to explore the developmental phasing of gene transcription programmes represented by C1-C3.

First, for single-cell RNA-seq data comprising 50,000 nuclei sampled from five cortical regions of three donor brains ^58^, the total weighted expression of the C1-C3 gene weights in each sample was computed separately for genes positively and negatively weighted in each component (**Methods**). We reasoned that if the components derived from bulk tissue microarray measurements in the AHBA dataset were merely reflective of regional differences in cellular composition, e.g. neuron-glia ratio, then genes weighted positively and negatively on each component should not have anti-correlated expression across cells of the same class. However, we observed that genes weighted positively and negatively on the same component had strongly anti-correlated expression at the single-cell level (**Fig. 3a**), whereas genes that were positively and negatively weighted on different components were not anti-correlated (**Supplementary Fig. S5**). The anti-correlation of genes positively and negatively weighted on C1 or C2 was stronger within each class of cells than across multiple cell classes; and even stronger when the single-cell data were stratified by sub-classes of cells in specific cortical layers, e.g., L2 VIP-expressing interneurons (**Fig. 3a inset**). In contrast, for C3 the anti-correlation of positively and negatively weighted genes was stronger across cell classes than within each class, although there was still evidence for significantly coupled expression across cells of the same class or subclass.

**Figure 3:**
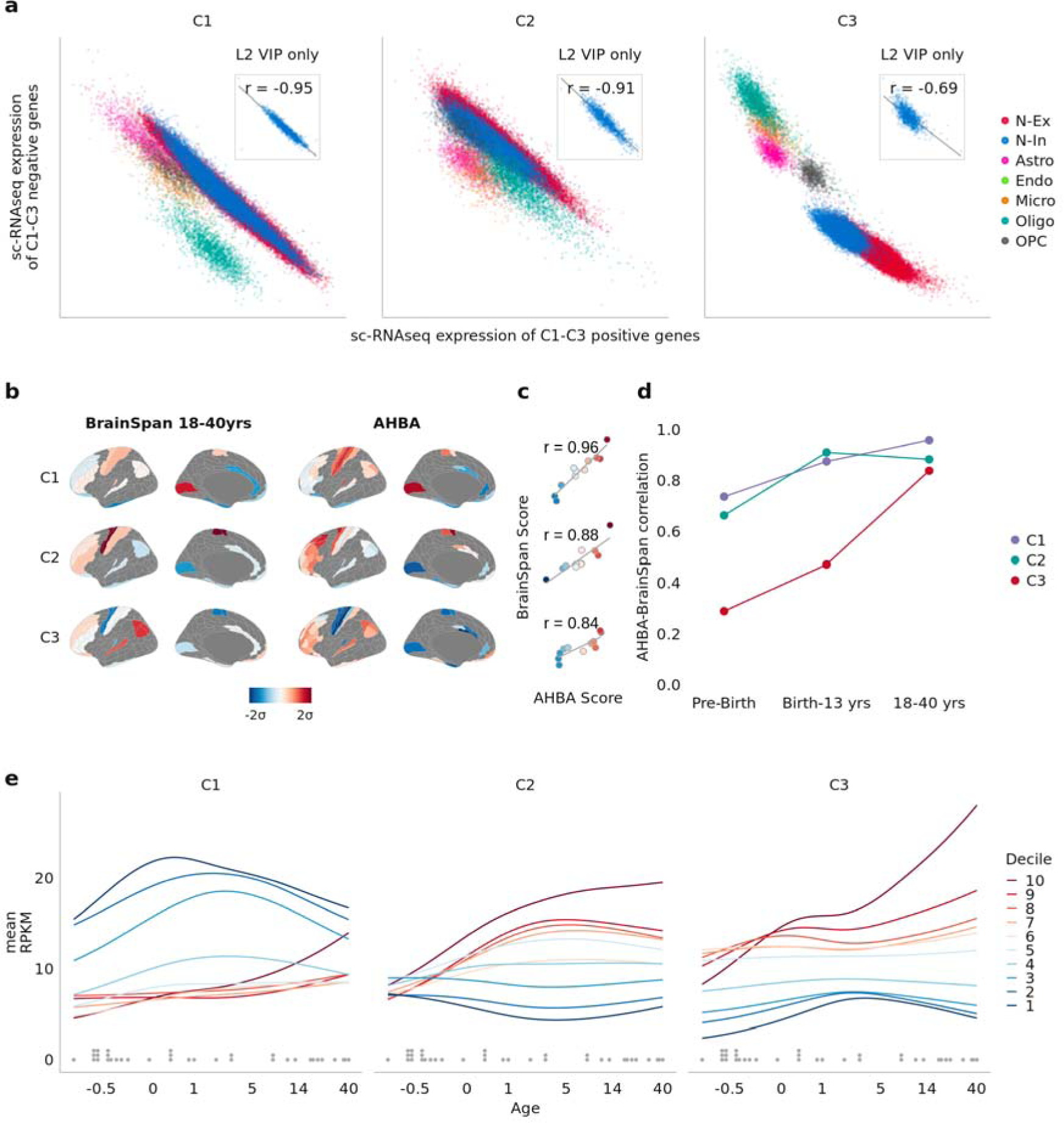
Transcriptional components represent intracellular coordination of gene expression programmes with distinct developmental trajectories. **a**, For each of ∼50,000 single-cell RNAseq samples, the weighted average expression of the negatively-weighted genes of each AHBA component C1-C3 is plotted against that of the positively-weighted genes (Methods). Samples are coloured by cell-type, demonstrating that genes positively and negatively weighted on C1-C3 have correlated expression within each major class of brain cells: N-Ex, excitatory neurons; N-In, inhibitory neurons; Astro, astrocytes; Endo, endothelial cells; Micro, microglia; Oligo, oligodendrocytes; and OPC, oligodendrocyte precursor cells. Inset, a subset of samples from Layer 2 VIP interneurons, illustrating that C1-C3 weighted genes were transcriptionally coupled even within a fine-grained, homogeneous group of cells. **b**, Cortical maps representing the regional scores of components C1-C3 for each of 11 regions with transcriptional data available in the BrainSpan cohort of adult brains (left) and C1-C3 component scores for the matching subset of regions in the AHBA (right). **c**, Scatter plots of matched regional C1-C3 scores from b, demonstrating that the three transcriptional components defined in the AHBA had consistent spatial expression in BrainSpan. **d**, Correlations between AHBA C1-C3 scores and BrainSpan C1-C3 scores (as in c) for each of 3 age-defined subsets of the BrainSpan dataset. C1 and C2 component scores were strongly correlated between datasets for all age subsets, whereas C3 component scores were strongly correlated between datasets only for the 18-40y subset of BrainSpan. This indicates that C1 and C2 components were expressed in nearly adult form from the earliest measured phases of brain development, whereas C3 was not expressed in adult form until after adolescence. **e**, Developmental trajectories of brain gene expression as a function of age (−0.5 to 40 years; x-axis, log scale) were estimated for each gene (Methods) and then averaged within each decile of gene weights for each of C1-C3; fitted lines are colour-coded by decile. Genes weighted positively on C3 were most strongly expressed during adolescence, whereas genes weighted strongly on C1 or C2 were most expressed in the first 5 years of life. Dots above the x-axis represent the post-mortem ages of the donor brains used to compute the curves. RPKM: reads per kilobase million.

Second, to explore the developmental trajectories of the transcriptional components, we used BrainSpan, an independent dataset where gene expression was measured by RNA-seq of bulk tissue samples from between 4 and 14 cortical regions for each of 35 donor brains ranging in age from −0.5 years (mid-gestation) to 40 postnatal years ^6^. We first asked if the gene weights for each of the components derived from the AHBA dataset would exhibit similar spatial patterns in the BrainSpan dataset. We projected the C1-C3 gene weights from the AHBA onto the subset of adult brains (18-40 years, N = 8) in BrainSpan (**Fig. 3b**, **Methods**) and found that the resulting cortical maps of component scores in the BrainSpan data were highly correlated with the corresponding cortical maps derived from the AHBA dataset (*r*_C1_ = 0.96, *r*_C2_ = 0.88, *r*_C3_ = 0.84; **Fig. 1d**). This indicated that the three components defined in the AHBA were generalisable to the adult brains in the BrainSpan dataset (for a full replication of C1-C3 in independent data see **Extended Data Fig. 3**). We then similarly compared the cortical component maps derived from the AHBA dataset to the corresponding maps calculated for subsets of the BrainSpan cohort from two earlier developmental stages (prebirth, N = 20, and birth-13 years, N = 14). We observed that for C1 and C2, AHBA component scores were almost as highly correlated with BrainSpan component scores in foetal (prebirth) and childhood (birth-13 years) brains as in the adult (18-40 years) brains (birth-13 years, *r*_C1_ = 0.87, *r*_C2_ = 0.91; prebirth, *r*_C1_ = 0.74, *r*_C2_ = 0.66; **Fig. 3d**). However, C3 scores in the AHBA dataset were not so strongly correlated with C3 scores in the foetal and childhood subsets of the BrainSpan dataset (prebirth, *r*_C3_ = 0.29; birth-13 years, *r*_C3_ = 0.47). These results suggested that C3 may only emerge developmentally during adolescence, whereas the C1 and C2 have nearly-adult expression from the first years of life.

We tested this hypothesis by further analysis of the BrainSpan dataset, modelling the non-linear developmental trajectories of each gene over the age range −0.5 to 40 years (**Methods**) and then averaging trajectories over all genes in each decile of the distributions of gene weights on each of the three components. We found that genes in the top few deciles of C3 gene weights became more strongly expressed during and after adolescence, whereas genes in the top few (C2) or bottom few (C1) deciles of gene weights on the other two components were most strongly expressed in the first 5 years of life and then declined or plateaued during adolescence and early adult life (**Fig. 3e**). These results confirmed that components C1-C3 have distinct neurodevelopmental trajectories, with genes positively weighted on C3 becoming strongly expressed after the first postnatal decade.

### Autism and schizophrenia have specific links to C1/C2 and C3

Finally, we explored the clinical relevance of C1-C3 by analysis of prior neuroimaging, differential gene expression, and GWAS associations for autism spectrum disorder (ASD), major depressive disorder (MDD), and schizophrenia.

First, we leveraged the BrainChart neuroimaging dataset of >125,000 MRI scans ^59^, in which atypical deviation of regional cortical volumes in psychiatric cases was quantified by centile scores relative to the median growth trajectories of normative brain development over the life-cycle (**Fig. 4a**). Using the Desikan-Killiany parcellation of 34 cortical regions necessitated by alignment with this dataset (**Methods**), we found that cortical shrinkage in ASD was significantly associated with both C1 and C2 (*r*_C1_ = 0.49, *p*_spin_ = 0.0002, FDR < 5%; *r*_C2_ = −0.28, *p*_spin_ = 0.0006, FDR < 5%), while shrinkage in schizophrenia was specifically associated with C3 (*r*_C3_ = 0.43, *p*_spin_ = 0.008, FDR < 5%) (**Fig. 4b**).

**Figure 4:**
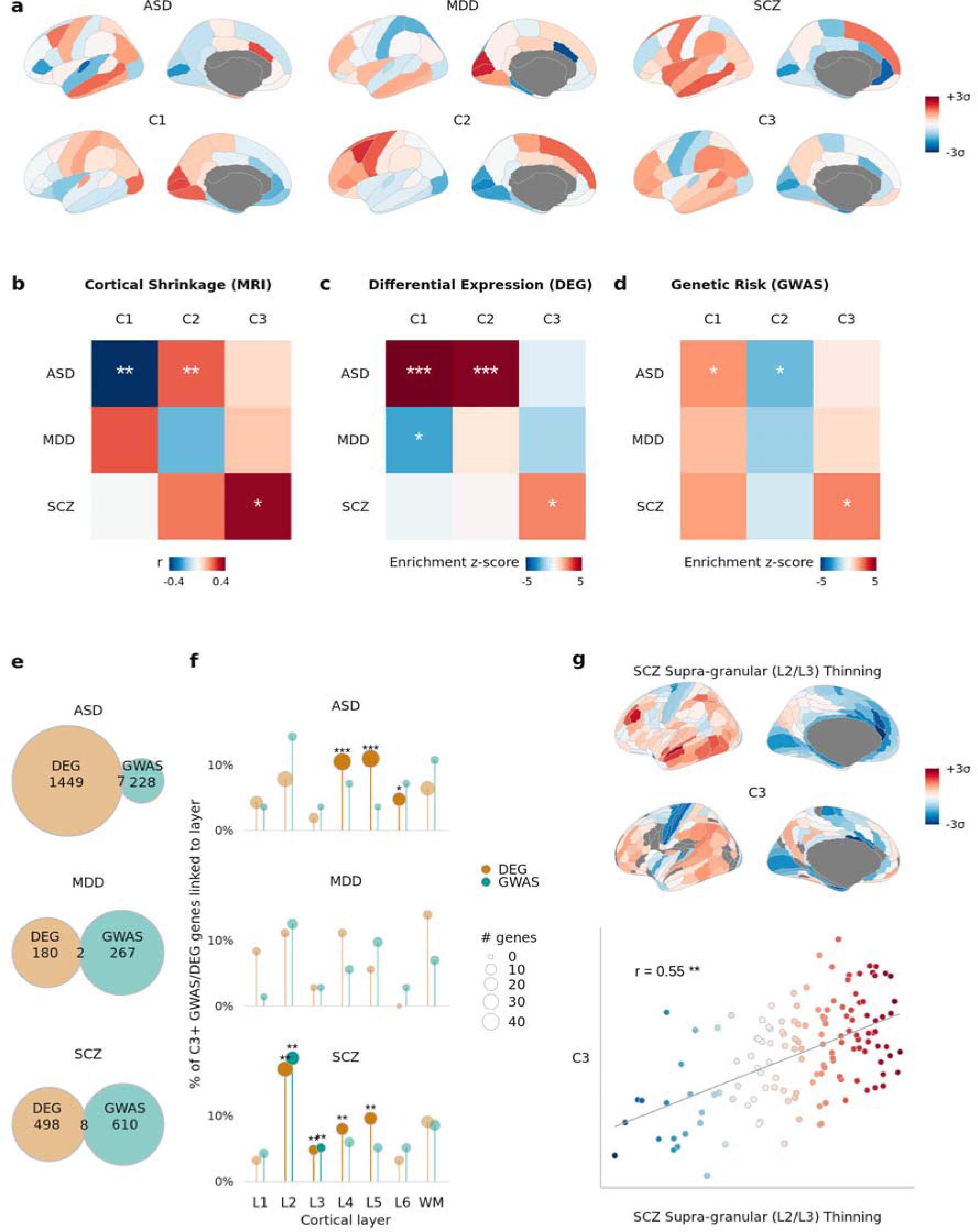
Genetics, transcriptomics, and neuroimaging of autism and schizophrenia were consistently and specifically linked to normative transcriptional programmes. **a**, First row: cortical volume shrinkage in autism spectrum disorder (ASD), major depressive disorder (MDD), and schizophrenia (SCZ) cases. Red indicates greater shrinkage, computed as z-scores of centiles from normative modelling of >125,000 MRI scans. Second row: AHBA components projected into the same Desikan-Killiany parcellation. **b**, Spatial correlations between volume changes and AHBA components, C1-C3. Significance tested by spatially autocorrelated spin permutations, and corrected for multiple comparisons: *, **, *** respectively indicate FDR-adjusted two-sided p-values: 0.05, 0.01, 0.001. **c**, Enrichments in C1-C3 for consensus lists of differentially expressed genes (DEGs) in postmortem brain tissue of donors with ASD, MDD, and SCZ compared to healthy controls (Methods). Significance assessed as percentile of mean weight of DEGs in each component relative to randomly permuted gene weights, and corrected for multiple comparisons: *, **, *** respectively indicate FDR-adjusted two-sided p-values: 0.05, 0.01, 0.001. **d**, Enrichment in C1-C3 for GWAS risk genes for ASD ^60^, MDD^61^, and SCZ ^62^, tested for significance as in c, demonstrating alignment with both spatial associations to volume changes and enrichments for DEGs. **e,** Venn diagrams showing the lack of overlap of DEGs and GWAS risk genes reported by the primary studies summarised in panels c and d. **f**, DEGs and GWAS risk genes for each disorder were filtered for only C3-positive genes, then tested for enrichment with marker genes for each cortical layer ^38^. Significance was tested by one-sided Fisher’s exact test and corrected for multiple comparisons across all 42 tests. C3-positive DEGs and GWAS genes for SCZ (but not ASD or MDD) were both enriched for L2 and L3 marker genes, despite the DEGs and GWAS gene sets having nearly no overlap for each disorder (see **Extended Data Fig. 6** for more detail). **g,** Convergent with L2/L3 enrichment in the C3-positive schizophrenia-associated DEGs and GWAS genes, a cortical map of supragranular-specific cortical thinning in schizophrenia ^63^ was significantly and specifically co-located with C3 (r = 0.55, two-sided spin-permutation p-value = 0.002); each point is a region, color represents C3 score.

Second, we compiled consensus lists of differentially expressed genes (DEGs) from RNA-seq measurements of dorsolateral prefrontal cortex tissue in independent studies of ASD ^36,64,65^, MDD ^66^, and schizophrenia ^65,67–70^ (**Methods**). We found that genes differentially expressed in ASD were specifically enriched in both C1 and C2 (but not C3); whereas genes differentially expressed in schizophrenia were enriched in C3 (but not C1 or C2); and genes differentially enriched in MDD were enriched only in C1 (**Fig. 4b**). Corroborating the enrichments of ASD DEGs, case-control differences in expression at 11 cortical regions for ASD cases compared to healthy controls showed the positively weighted genes on C1 and C2 were significantly less strongly expressed in ASD cases than in controls (**Extended Data Fig. 3**).

Third, using data from the most recent GWAS studies of ASD ^60^, MDD ^61^, and schizophrenia ^62^, we found that genetic variants significantly associated with ASD were enriched in both C1 and C2 (but not C3); whereas genes associated with schizophrenia were enriched in C3 (but not C1 or C2) (**Fig. 4d**). Genes associated with MDD were not significantly enriched in any transcriptional component. These associations were replicated when using alternative methods (MAGMA ^71^ and H-MAGMA ^72^) to test the association between GWAS-derived p-values for the association of each gene with ASD, MDD or schizophrenia and the C1-C3 gene weights without requiring an explicit prioritisation of GWAS-associated genes (**Supplementary Fig. S6**). This pattern of results for autism and schizophrenia GWAS associations evidently mirrored the pattern of prior results from analysis of case-control neuroimaging (**Fig. 4b**) and differential gene expression studies (**Fig. 4c**), with ASD consistently linked to components C1 and C2, and schizophrenia consistently linked to C3.

Notably, this consistency of association between disorders and specific transcriptional components was observed despite minimal overlap between the DEGs and GWAS risk genes identified as significant by the primary studies of each disorder ^73^ (**Fig. 4e**). However, motivated by the association of C3 with regions of greatest laminar differentiation (**Fig. 2b**), we found that the subsets of the schizophrenia-associated DEG and GWAS gene sets that were positively-weighted on C3 were both significantly enriched for marker genes of layers L2 and L3 (**Fig. 4g; Extended Data Fig. 6**). These shared laminar associations between the non-overlapping DEG and GWAS gene sets were only present when subsetting to C3-positive genes, and were specific to schizophrenia (i.e. C3-positive subsets of ASD and MDD genes did not show the same L2/L3 enrichments). Convergent with C3 revealing an L2/L3 association in schizophrenia-associated genes from DEG and GWAS, we found that the cortical map of C3 was significantly co-located with an MRI-derived map of specifically supragranular, L2/L3 predominant thinning in schizophrenia ^63^ (*r* = 0.55, *p*_spin_ = 0.002, FDR < 1%, **Fig. 4g**).

## Discussion

Our results offer a new perspective on how the brain’s macroscale organisation develops from the microscale transcription of the human genome. Through optimized processing of the AHBA and replication in PsychENCODE, we have shown that the transcriptional architecture of the human cortex comprises at least three generalisable components of coordinated gene expression. The two higher-order components (C2 and C3) had not previously been robustly demonstrated, although the initial AHBA paper identified similar components to C1 and C2 by applying PCA to one of the six AHBA brains and filtering for only 1000 genes ^2^ (**Supplementary Fig. S1**). Here we derive C2 and C3 from all six AHBA brains and show they each represent the coordinated expression of hundreds of genes (**Supplementary Fig. S2**). Broadly, the C2 genes were enriched for “metabolic” and “epigenetic” processes, while the C3 genes were enriched for “synaptic” and “immune” processes (**Fig. 1c**). Both higher-order components were significantly enriched for genes associated with intelligence and educational attainment (**Fig. 1f-g**), indicating their relevance to the brain’s ultimate purpose of generating adaptive behaviour. The brain maps corresponding to each of the components were also distinctively co-located with multiple neuroimaging or other macroscale brain phenotypes (**Fig. 2**). These co-locations were often convergent with the gene enrichment results, triangulating evidence for C2 as a metabolically specialised component and for C3 as a component specialised for synaptic and immune processes underpinning adolescent plasticity; see **Table 1**. Together, these convergent results expand on the proposal of a single “sensorimotor-association axis” ^10,74^ by demonstrating that macro-scale brain organisation emerges from multiple biologically-relevant transcriptional components.

**Table 1:** Summary of convergent results on the biological and clinical relevance of three human brain transcriptional programmes. Each of three components of normative human brain gene expression (C1, C2, C3; table columns) was biologically validated by testing for enrichment of gene weights on each component, and for co-location of regional component scores with neuroimaging or other macro-scale brain phenotypes, in healthy brain samples (normative) and in studies of neurodevelopmental disorders (atypical). Each row summarises results for a distinct gene enrichment analysis (italicised) or spatial co-location analysis (plain font). Based on prior knowledge that theta oscillations are linked to intelligence and cognition ^75^ as well as to glucose metabolism ^76^, the spatial alignments between C2 and maps of MEG theta power (**Fig. 2d**) and aerobic glycolysis (**Fig. 2a**) were convergent with the enrichment of C2 for genes linked to cognitive capacity (**Fig. 1f-g**) and metabolism (**Fig. 1c**). Similarly, prior knowledge implicates microglia and oligodendrocytes in the immune-mediated synaptic pruning and myelination that over adolescence gives rise to adult cognitive capacity ^77,78^, such that the spatial alignment between C3 and the map of adolescent myelination (**Fig. 2d**) was convergent with the enrichments of C3 for genes related to immunity, synaptic development, and learning (**Fig. 1c**); oligodendrocytes, microglia, and synapses (**Fig. 1e**); and cognitive capacity (**Fig. 1f-g**), among which one GWAS study explicitly linked intelligence to myelination ^41^.

|  |  | **C1: Neuronal hierarchy** | **C2: Cognitive metabolism** | **C3: Adolescent plasticity** |
| --- | --- | --- | --- | --- |
| **Normative** | **Biological processes**<br>(Fig. 1c) | *Most genes are aligned, especially PVALB, SST* | *Metabolism*<br>*Epigenetics* | *Synaptic plasticity*<br>*Learning/memory*<br>*Immunity* |
|  | **Architectonics**<br>(Fig. 1d) | *L4*<br>*L1, L2, L6* | *L4, L5, L6*<br>*L2* | *L2, L3, L4, L5, L6*<br>*L1, WM* |
|  | **Cell types**<br>(Fig. 1e) | *Oligodendrocytes*<br>*Astrocytes* | *Synapses,*<br>*Endothelial cells* | *Synapses, Neurons*<br>*Oligodendrocytes,*<br>*Microglia* |
|  | **GWAS**<br>(Fig. 1f-g) | *Educational attainment* | *Intelligence/cognition*<br>*Educational attainment* | *Intelligence/cognition*<br>*Educational attainment* |
|  | **Imaging**<br>(Fig. 2) | *fMRI degree*<br>*T1w/T2w*<br>*Cortical thickness* | *MEG theta power*<br>*Aerobic glycolysis* | *Adolescent change in myelination* |
|  | **Development**<br>(Fig. 3b-c) | *Prenatal, greatest expression at birth* | *Prenatal, greatest expression in first decade* | *Adolescence, greatest expression in adulthood* |
| **Atypical** | **Imaging**<br>(Fig. 4a-b, g) | ASD volume shrinkage | ASD volume shrinkage | SCZ volume shrinkage and L2, L3-specific thinning |
|  | **RNA-seq of brain tissue**<br>(Fig. 4c,f) | ASD DEGs | ASD DEGs | SCZ DEGs, with L2, L3 enrichment |
|  | **GWAS**<br>(Fig. 4d,f) | ASD risk genes | ASD risk genes | SCZ risk genes, with L2, L3 enrichment |

The discovery of these biologically-relevant, higher-order transcriptional components in the AHBA dataset raised further questions: i) do the components reflect coordinated gene expression within cells, or only variation in cell composition; ii) when do the components emerge during brain development; and (iii) how do they intersect with neurodevelopmental disorders? We addressed these questions using additional RNA-seq datasets (**Supplementary Table S5**). First, we found that genes positively or negatively weighted on the components derived from the AHBA bulk tissue samples had consistently coupled co-expression across RNA-seq measurements in single cells, e.g. individual neurons and glia (**Fig. 3a**). This indicated that C1-C3 represent transcriptional programmes coordinated at the intracellular level, not merely regional variation in the proportion of different cell types. Second, we found that C1-C3 have differentially phased developmental trajectories of expression, e.g. that the positive pole of C3 becomes strongly expressed only during adolescence, convergent with its spatial co-location with a map of adolescent cortical myelination (**Fig. 3b-c**). Finally, we established that these transcriptional programmes are not only critical for healthy brain development but, as might be expected, are also implicated in the pathogenesis of neurodevelopmental disorders (**Fig. 4**).

The pattern of results for disorders was strikingly convergent across multiple data modalities: C1 and C2 were both enriched for genes implicated by both GWAS and DEG data on ASD, whereas C3 was specifically enriched for genes implicated by both GWAS and DEG data on schizophrenia (**Table 1**). We observed a similar pattern of significant co-location between C1-C3 maps and MRI phenotypes: developmentally normalised scores on reduced cortical volume in ASD were correlated with maps of C1 and C2, and for schizophrenia with the map of C3 (**Fig. 4a-b**). In contrast, there was no evidence for enrichment of C1-C3 by genes associated with risk of Alzheimer’s disease ^79^ (**Supplementary Fig. S6**). An intuitive generalisation of these results is that the developmental processes which give rise to these three components of gene expression in the healthy adult brain are pathogenically more relevant for neurodevelopmental disorders than for neurodegenerative disease.

Overall, our results were strongly supportive of the motivating hypothesis that the transcriptional architecture of the human cortex represents developmental programmes crucial both to the brain’s healthy organisation and to the emergence of neurodevelopmental disorders. For example, when interpreting C3 as a transcriptional programme mediating adolescent plasticity (**Table 1**), our finding that C3 represents coupled transcription of synapse- and immune-related genes within cells (**Fig. 3a**) is consistent with prior work indicating that the neuronal expression of immune-related, typically glial genes can play a mechanistic role in synaptic pruning ^80^; and, vice-versa, that neuronal genes associated with synapse and circuit development can also be expressed in glial cells ^81^. While atypical synaptic pruning has long been hypothesised to be a mechanistic cause of schizophrenia ^82–84^, prior results on the biology of schizophrenia have shown limited consistency, both between the primary data modalities of GWAS, post-mortem expression, and neuroimaging ^85,86^, and even between DEG studies ^73^. Here, we demonstrate that the C3 transcriptional programme offers a unifying link between these disparate prior results. When parsed by the C3 positive genes, the otherwise non-overlapping GWAS and DEG gene-sets for schizophrenia display a shared enrichment for supra-granular marker genes (**Fig. 4e-f**), and, convergently, C3 was spatially associated with supra-granular-specific thinning in schizophrenia (**Fig. 4g**). Supra-granular layers have dense cortico-cortical connections and are expanded in humans relative to other species ^87–89^, mature latest in development ^90^, have been linked to intelligence ^91^, and have previously been linked to schizophrenia ^92–94^. This triangulation of evidence strongly suggests that the third component of the brain’s gene expression architecture represents the transcriptional programme coordinating the normative, neuro-immune processes of synaptic pruning and myelination in adolescence ^56^, such that atypical expression of C3 genes due to schizophrenia genetic risk variants can result in atypical development of supra-granular cortical connectivity leading to the clinical emergence of schizophrenia.

Clearly there are limits to what can be learnt from RNA measurements of bulk tissue samples from six healthy adult brains. Here, we explicitly identified the limits of the AHBA dataset by optimizing data processing against an unbiased measure of generalisability, *g*, which yielded three components. The architecture of human brain gene expression likely involves more than three components; however, our analysis suggests that their discovery will rely on additional high-granularity transcriptional data. In particular, gene expression varies with sex, age, genetics, and environment ^95^, so we expect that future data will reveal additional components that are more individually variable and demographically diverse than the three we have characterised here. Meanwhile, the code and data that supplement our results can help future research to leverage our work with the unique AHBA resource.

## Extended Data Figures

**Extended Data Fig. 1:**
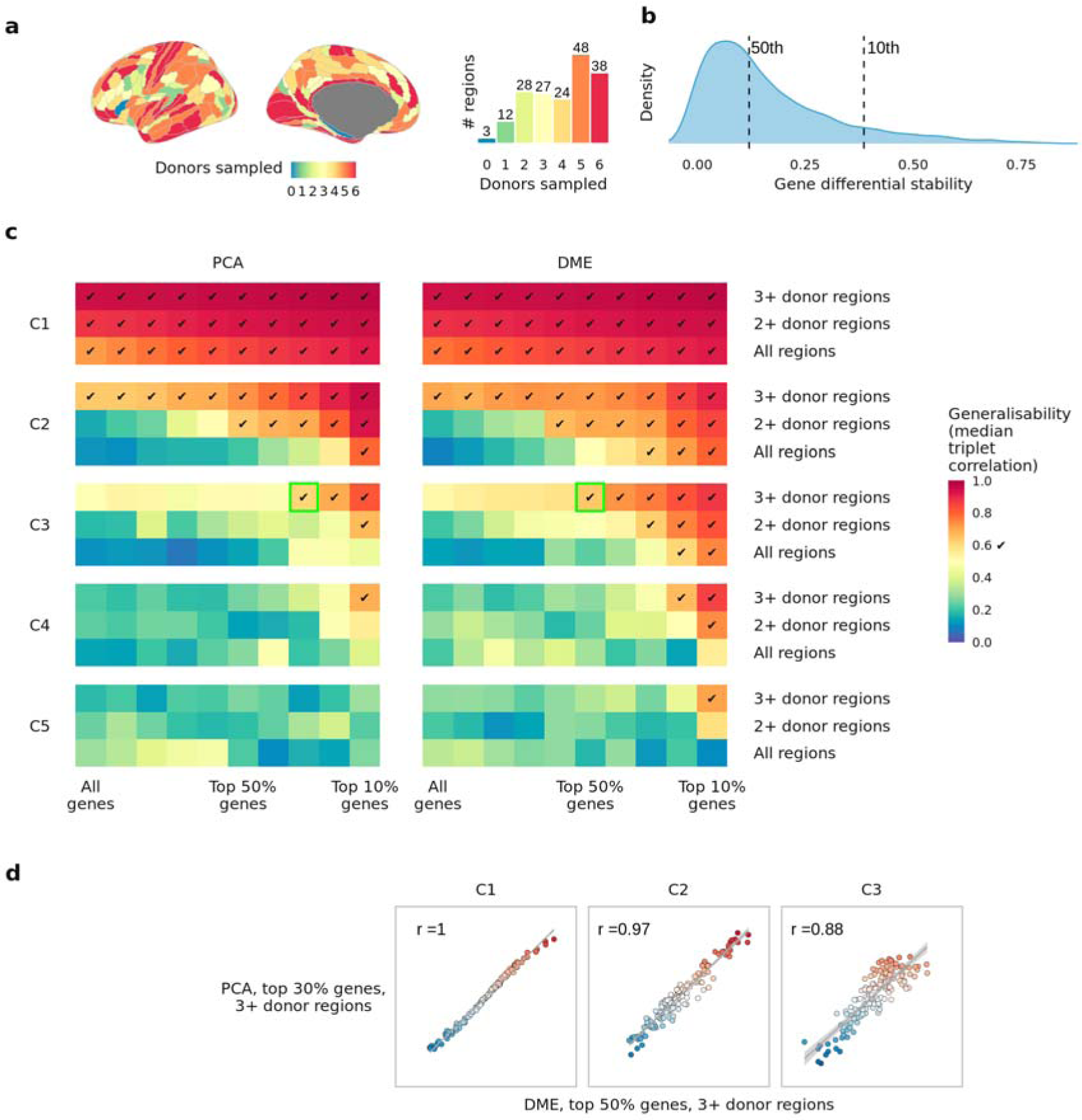
Optimised processing of the AHBA identified three generalisable components. **a**, In the HCP-MMP parcellation, 43/180 regions are matched to samples representing less than 3 of the 6 AHBA donors.**b**, Distribution of differential stability of genes measured in the AHBA dataset processed in the HCP-MMP parcellation. **c**, Generalisability of first five components of the AHBA dataset computed with either principal components analysis (PCA) or diffusion map embedding (DME). Color represents generalisability g, defined as the median absolute correlation between matched components computed across all 10 disjoint triplet pairs (Methods); x-axis represents variation in the proportion of genes filtered out by differential stability prior to PCA/DME; y-axis represents variation in which regions are filtered out prior to PCA/DME. Tick mark indicates parameter combinations that exceed generalisability g > 0.6. Green highlights for C3 indicate the best parameter option with PCA and DME respectively, showing that switching to DME achieves similar generalisability while retaining more genes. **d**, Scatter plots of regional scores for AHBA components computed using the best PCA/DME options, demonstrating that PCA and DME derive spatially equivalent components.

**Extended Data Fig. 2:**
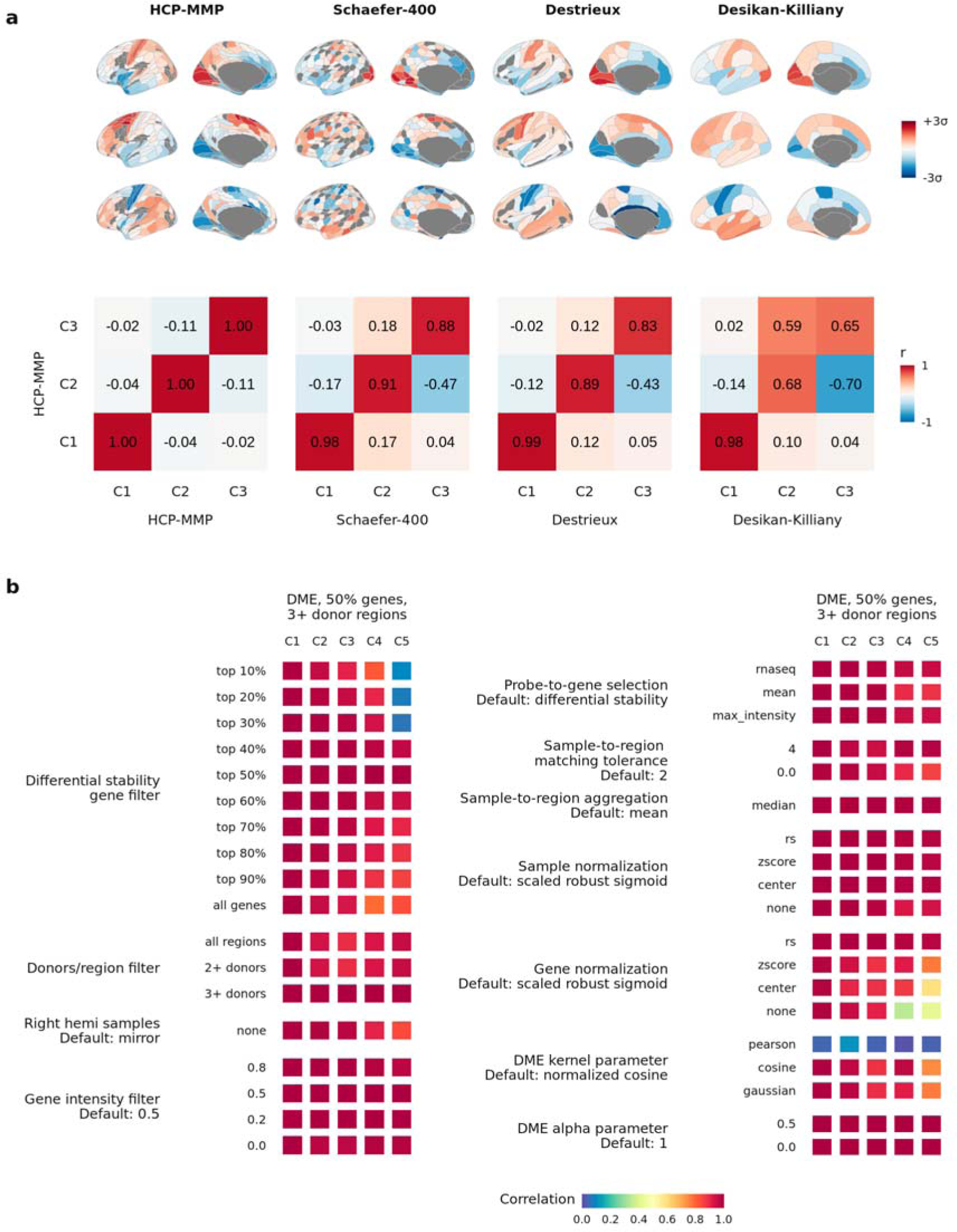
Transcriptional components were robust to parcellation and processing. Transcriptional components were computed in four different parcellation templates (Methods). For each parcellation, the gene weights for the first three components were correlated with the weights obtained from the HCP-MMP parcellation used throughout. Gene weights were highly consistent, although in the less-granular (34-regions/hemisphere) Desikan-Killiany parcellation, C2 and C3 were less well aligned to the other parcellations. **b**, A wide range of parameters for processing the AHBA data were varied, and the resulting component region scores were correlated with the components obtained from the optimised parameters. For nearly all variations in parameters, highly consistent components were obtained, demonstrating the robustness of C1-C3.

**Extended Data Figure 3:**
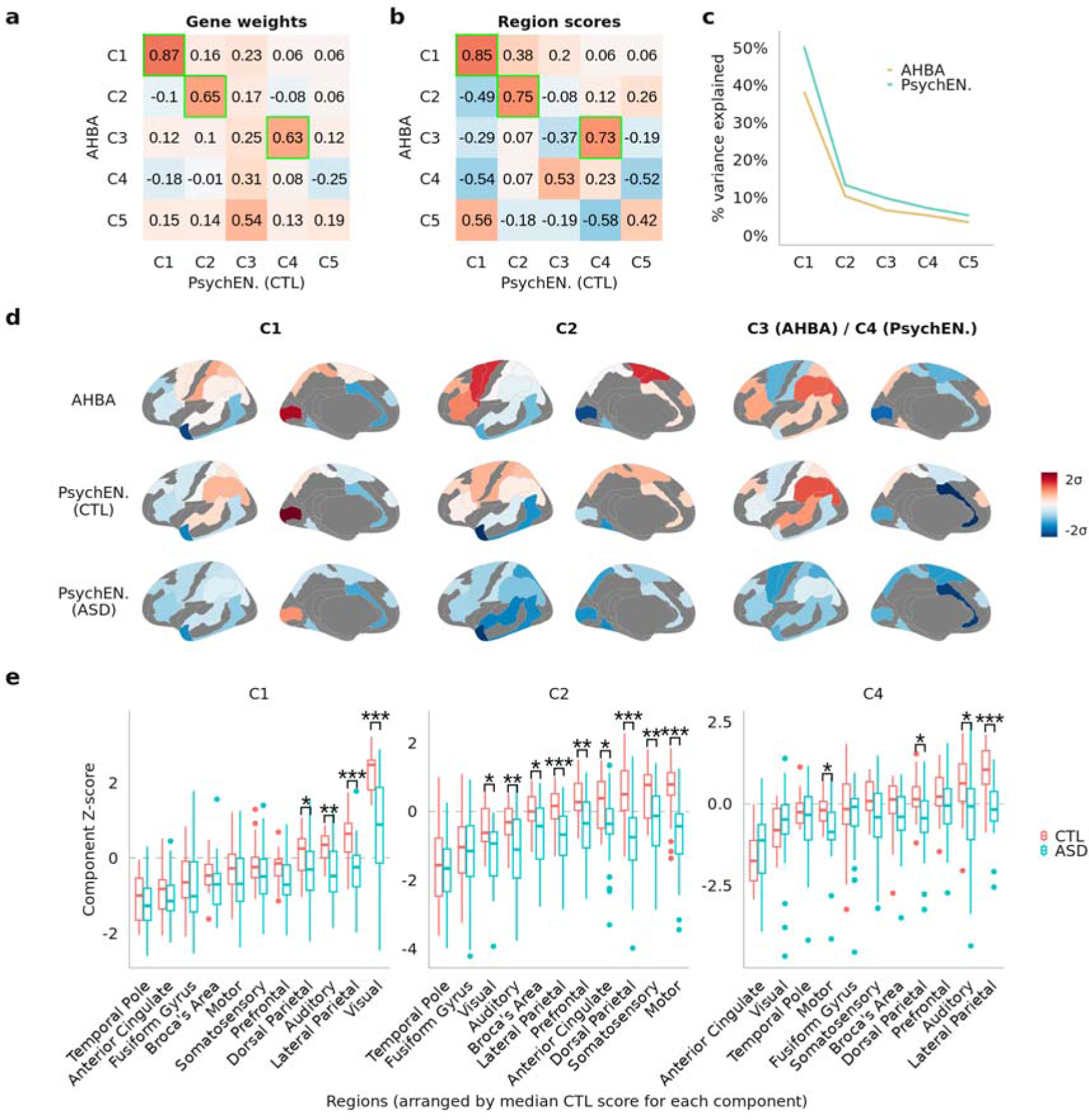
AHBA transcriptional components were reproducible in independent PsychENCODE control data, with differential spatial expression in autism. **a**, Gene weights from dimension reduction applied to group-averaged bulk RNA-seq measurements from 11 cortical regions in N = 54 healthy control brains from the PsychENCODE dataset ^36^ were correlated with gene weights from the components of the AHBA (derived by DME in the 180-region HCP-MMP parcellation), showing that the genetic profiles of AHBA C1, C2, and C3 were reproduced by PsychENCODE C1, C2, and C4, respectively (highlighted in green). **b**, Regional scores of PsychENCODE C1, C2 and C4 were also correlated with region scores of AHBA C1, C2 and C3, showing that the matching genetic profiles correspond to matching spatial expression patterns. **c**, Variance explained by the first five components of each dataset, showing that AHBA C3 and PsychENCODE C4 account for similar proportions of variance (6.5% and 7.1%, respectively). **d**, 1st row: Cortical maps of AHBA C1-C3 in the same 11 regions sampled in the PsychENCODE data. 2nd row: Cortical maps of PsychENCODE C1, C2, and C4 demonstrating their spatial similarity to AHBA C1-C3. 3rd row: Gene weights from the PsychENCODE healthy control data were projected onto transcriptional data of cases with autism spectrum disorder (ASD; N = 58) from the same dataset, demonstrating lower regional expression at the positive(red) pole of each component in the ASD cases compared to healthy controls. **e**, Distributions of regional scores for C1, C2 and C4, computed on group-average healthy controls as in a-d and projected to individual donor brains in the PsychENCODE dataset, demonstrating significant case-control differential expression for regions at the positive poles of C1-C3. T-tests of case-control differences were corrected for multiple comparisons across all 33 tests; boxplots represent the median, first, and third quartiles with whiskers showing 1.5 * inter-quartile range; *, **, *** indicate FDR-corrected two-sided p-value < 0.05, 0.01, 0.001 respectively. Region names refer to the sampled Brodmann Areas (BA) ^36^: Visual = BA17, Temporal Pole = BA38, Somatosensory = BA3-1-2-5, Motor = BA4-6, Anterior Cingulate = BA24, Prefrontal = BA9, Broca’s Area = BA44-45, Fusiform Gyrus = BA20-37, Auditory = BA41-42-22, Lateral Parietal = BA39-40, Dorsal Parietal = BA7.

**Extended Data Figure 4:**
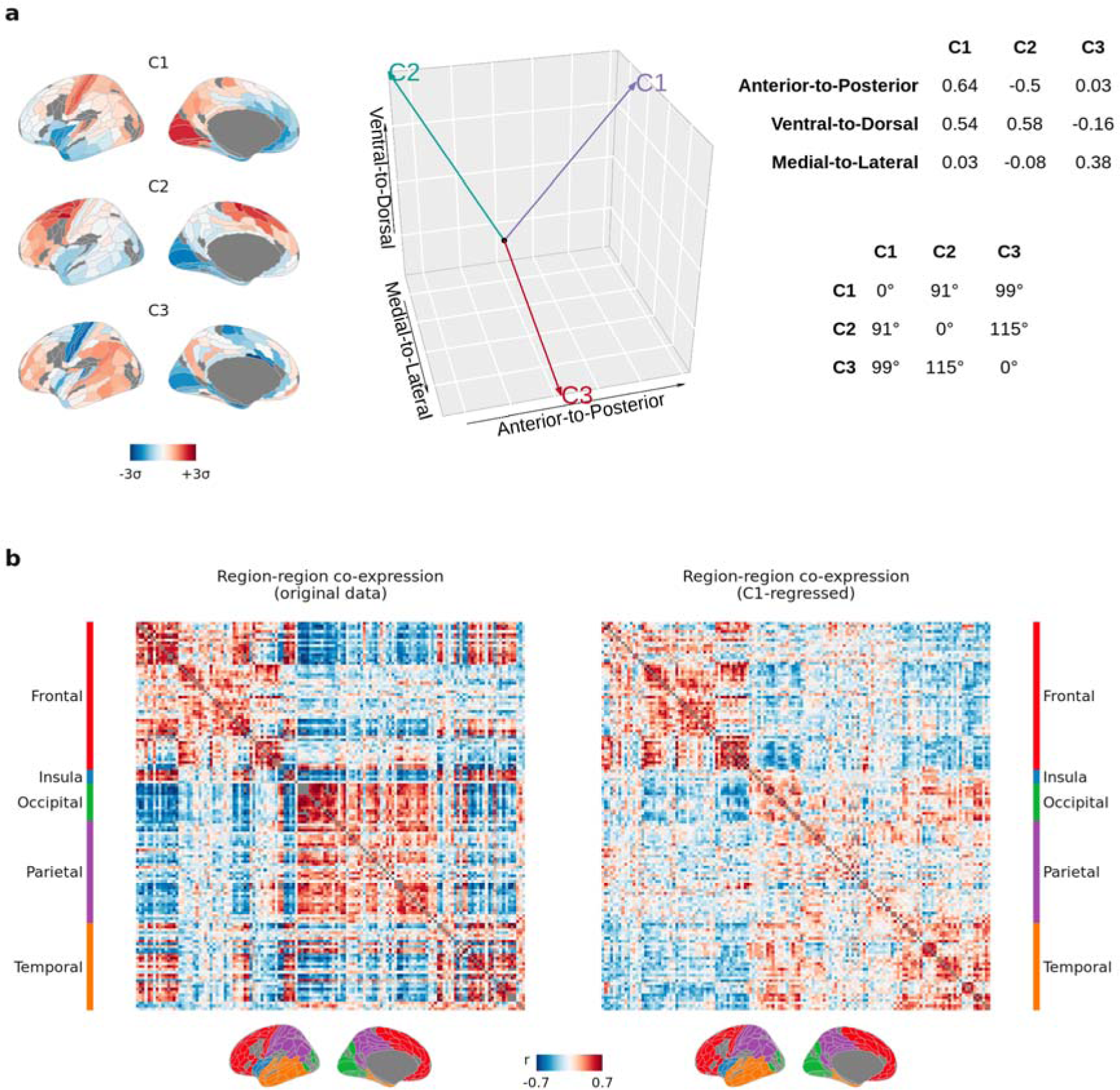
Higher-order components of cortical gene expression reflect anatomically relevant co-expression structure. **a,** C1-C3 were orthogonally aligned in anatomical space, as computed by the Pearson’s correlations of the regional scores with the XYZ coordinates of the region centroids: C1 and C2 were both aligned with the anterior-to-posterior (y) and ventral-to-dorsal (z) plane, but with opposite signs along the anterior-to-posterior axis, while only C3 was aligned to the medial-lateral (x) axis. The middle panel represents these alignments as vectors in 3D space. The right-hand upper table shows the correlations of C1-C3 with each anatomical axis, and the lower table shows the angle in degrees between the vectors, showing that C1-C3 are orthogonal. **b**, Co-expression matrices computed by Pearson’s correlations of gene expression between brain regions, computed with and without regressing out the first component C1, and annotated by the major cortical lobes as defined in the HCP-MMP parcellation ^32^. This further demonstrates that the gene co-expression structure captured by C2 and C3 (i.e., the residual variation beyond C1) is anatomically relevant.

**Extended Data Figure 5:**
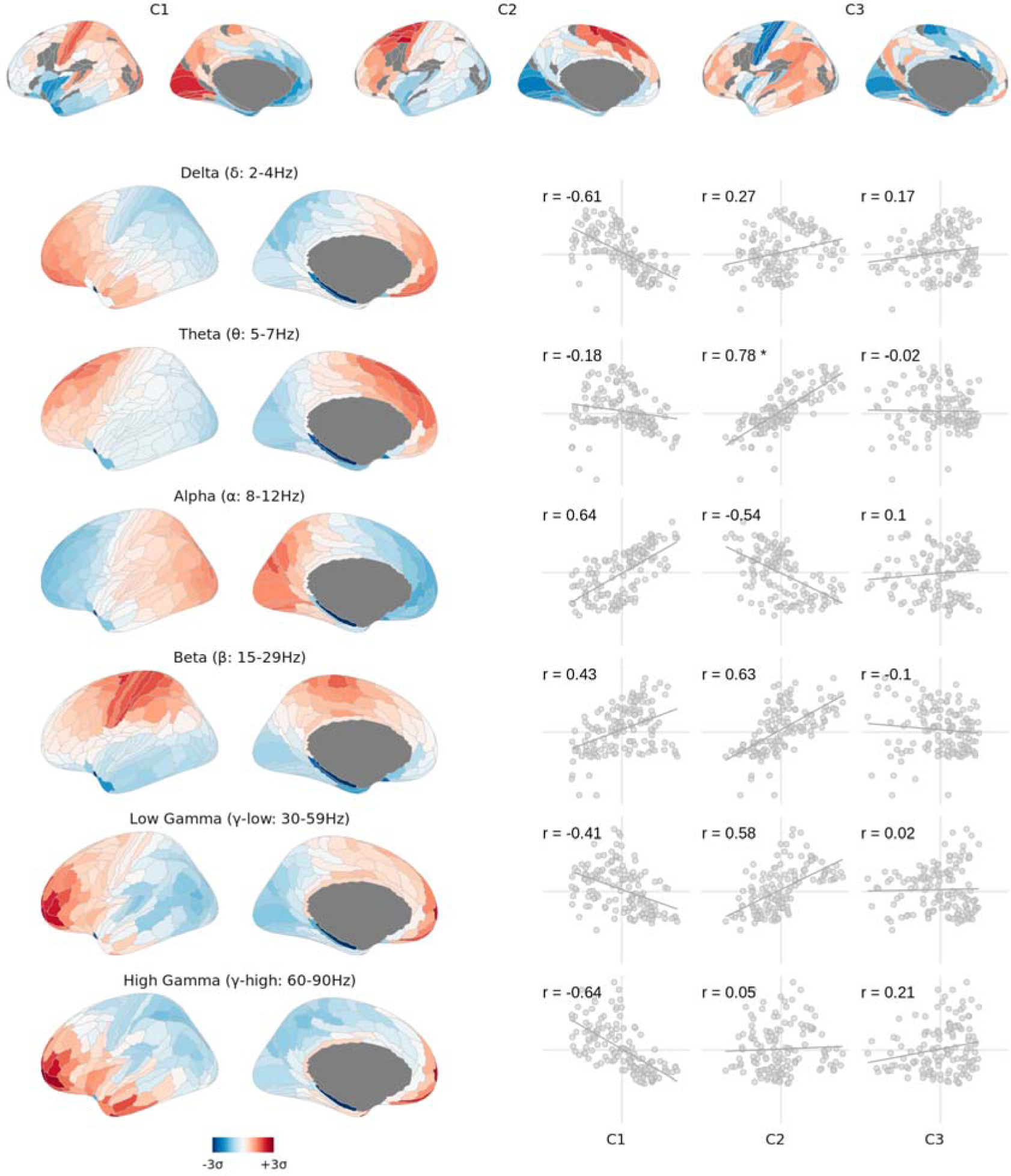
Transcriptional components were distinctively associated with the regional power of canonical brain oscillation frequencies. Several MEG power bands ^55^ were highly correlated (|r|>0.6) with C1 (delta, alpha, high-gamma) and C2 (beta, theta), although only the theta association to C2 survived FDR correction of the spin-test p-values (r=0.78, FDR_spin_=0.05). No MEG band was aligned with C3.

**Extended Data Figure 6:**
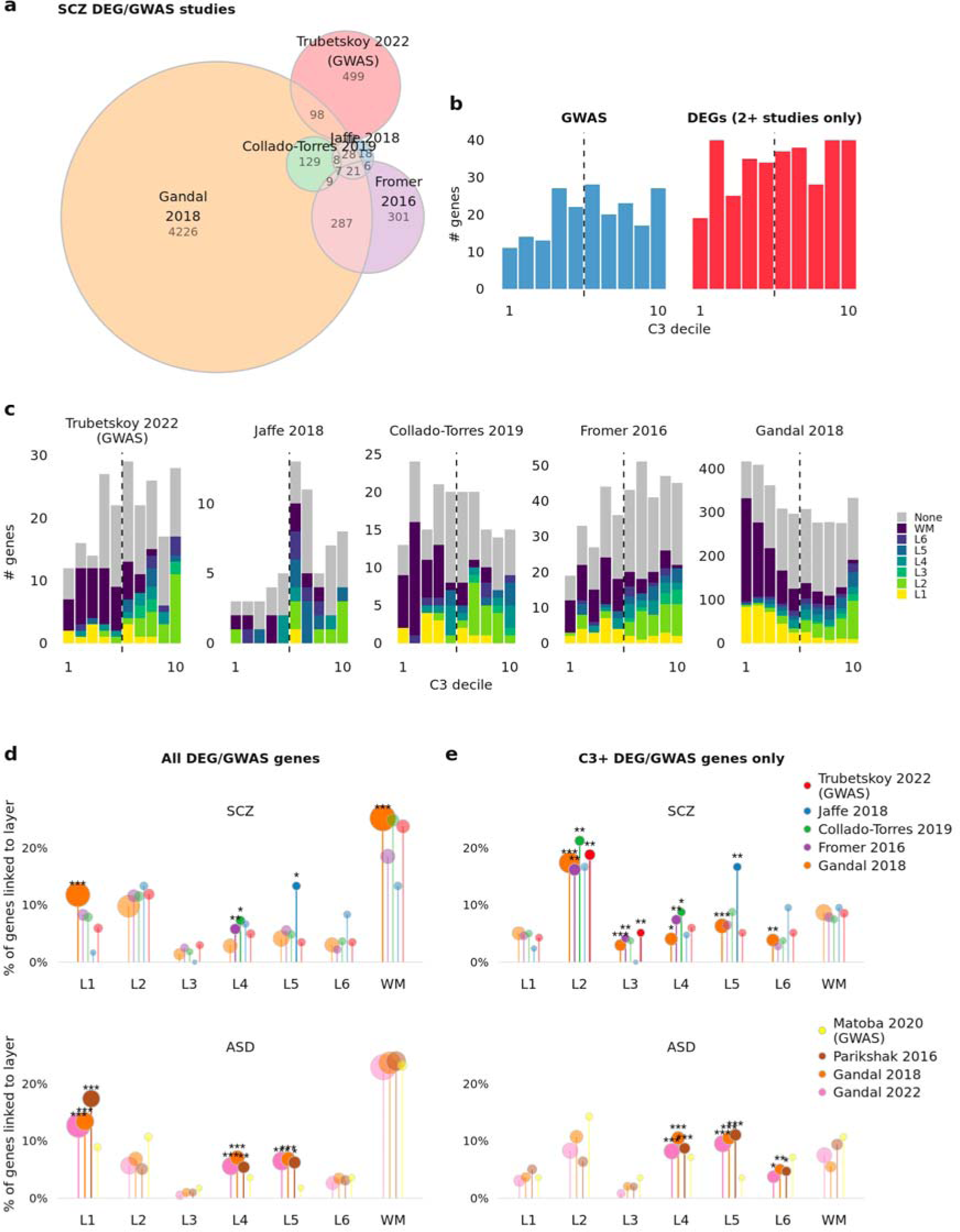
C3 reveals shared biology across inconsistent postmortem brain RNA-seq studies of differentially expressed genes (DEGs) in schizophrenia. **a**, Euler diagram demonstrating the relative lack of overlap of genes linked to schizophrenia in four independent RNA-seq postmortem brain studies, as well as the latest GWAS study. **b**, Histogram of the schizophrenia GWAS and consensus DEG genes by C3 decile. The skew of the histograms towards higher C3 deciles reflects the significant enrichment of both non-overlapping gene sets, as in Fig. 4c-d. **c**, Histograms of the schizophrenia GWAS and DEG genes from each separate study by C3 decile, coloured by cortical layer where the gene was identified as a marker gene ^38^. L2 genes are distinctly clustered towards the C3+ pole, while L1 and WM genes are clustered towards C3-. **d**, For schizophrenia and ASD, enrichments of the GWAS/DEG genes from each separate study for marker genes of cortical layers, showing that no consistent significant enrichments are found across t he entire gene sets for studies of either disorder. **e**, Enrichments as in d, except for only genes positively weighted in C3 (corresponding to the right-hand five deciles of each histogram in panel c). For schizophrenia, significant enrichments for L2 and L3 are observed for three of the four DEG studies, as well as the GWAS study. No such enrichments were observed for ASD, demonstrating that C3 reveals convergent biology across otherwise inconsistent results specifically for schizophrenia. Significance was tested by one-sided Fisher’s exact test and corrected for multiple comparisons across all tests in each panel. *, **, *** indicate FDR-corrected one-sided p-value < 0.05, 0.01, 0.001 respectively.

## Acknowledgements

The authors acknowledge and thank Sofie Valk and Varun Warrier for their helpful comments on the manuscript.

R.D. was supported by the Gates Cambridge Trust. J.S. was supported by NIMH T32MH019112-29 and K08MH120564. A.A. was funded by a grant from the Australian Research Council (ARC) under its Linkage Project scheme (LP160101592). R.A.I.B. was supported by the Autism Research Trust. K.S.W. was supported by the Wellcome Trust (215901/Z/19/Z). E.T.B. was supported by an NIHR Senior Investigator award and the Wellcome Trust collaborative award for the Neuroscience in Psychiatry Network (NSPN). A.R. was supported by the National Institute of Mental Health Intramural Research Program (NIH Annual Report Number, 1ZIAMH002949-04). P.E.V. is a Fellow of MQ: Transforming Mental Health (MQF17_24). The funders had no role in study design, data collection and analysis, decision to publish or preparation of the manuscript.

Data were curated and analysed using a computational facility funded by an MRC research infrastructure award (MR/M009041/1) to the School of Clinical Medicine, University of Cambridge. All research at the Department of Psychiatry in the University of Cambridge is supported by the NIHR Cambridge Biomedical Research Centre (BRC-1215-20014) and NIHR Applied Research Centre. The views expressed are those of the authors and not necessarily those of the NIH, NHS, the NIHR or the Department of Health and Social Care. For the purpose of open access, the authors have applied a Creative Commons Attribution (CC BY) licence to any Author Accepted Manuscript version arising from this submission.

## Author Contributions statement

P.E.V., R.D., J.S., K.W., and A.R. designed research. J.S., A.A., K.W., and R.B. contributed data. R.D. and P.E.V. performed research, and E.T.B., K.A., K.W., A.R., J.S., R.M., and A.A. helped interpret results. R.D., P.E.V., and E.T.B. wrote the manuscript. All authors reviewed the manuscript.

## Methods

### AHBA data and donor-level parcellation images

Probe-level gene expression data with associated spatial coordinates were obtained from the Allen Institute website (https://human.brain-map.org), which collected the data after obtaining informed consent from the deceased’s next-of-kin. HCP-MMP1.0 parcellation images matched to the individual native MRI space of each donor brain (N=6) were obtained from Arnatkevičiūtė *et al.* (https://figshare.com/articles/dataset/AHBAdata/6852911) ^33^. The use of native donor parcellation images (rather than a standard parcellation image with sample coordinates mapped to MNI space) was chosen as it optimised the triplet generalisability metric (see following).

### AHBA processing parameters

To correctly match AHBA samples to regions in native donor space parcellation images using published processing pipelines, we recommend the use of either (i) *abagen* version 0.1.3 or greater (for Python) ^34^, or (ii) the version of the *AHBAprocessing* pipeline updated in June 2021 or later (for Matlab) ^33^.

Here we processed the AHBA with the *abagen* package, with one modification: we filtered the AHBA samples for only those annotated as cortical samples prior to subsequent processing steps. This was done such that subcortical and brainstem samples did not influence the intensity filter and probe aggregation steps. This modification was chosen as it optimised the triplet generalisability metric (see following). The code used to apply the modification is available in the *code/processing_helpers.py* file at https://github.com/richardajdear/AHBA_gradients.

Other than this modification, *abagen* was run using the following parameters, which follow published recommendations ^33^ unless otherwise specified:

- *Hemisphere*: The right hemisphere samples that are present for two of the six donors were reflected along the midline and processed together with the left hemisphere samples of those donor datasets to increase sample coverage.
- *Intensity-based filter* : Probes were filtered to retain only those exceeding background noise (as defined by the binary flag provided with the data by the Allen Institute) in at least 50% of the samples ^33^.
- *Probe aggregation*: Probes were aggregated to genes by differential stability, meaning that for each gene, the probe with the highest mean correlation across donor pairs was used.
- *Distance threshold* : Samples were matched to regions with a tolerance threshold of 2mm, using the voxel-mass algorithm in the *abagen* package.
- *Sample normalisation*: Prior to aggregating over donors, samples were normalised across all genes, using the scaled robust sigmoid method (a sigmoid transformation that is robust to outliers ^33^).
- *Gene normalisation* : Prior to aggregating over donors, genes were normalised across all samples, again using the scaled robust sigmoid.

To ensure robustness, the primary analysis of computing components of the AHBA was repeated in a series of sensitivity analyses varying all of the processing parameters above, e.g. not mirroring right hemisphere samples to the left hemisphere, different or no intensity filter for genes, different methods for aggregating and normalising probes. Sensitivity analyses also included running the pipeline with alternative parcellation templates: HCP-MMP1.0 ^32^, Schaefer-400 ^96^, Desterieux ^97^, and Desikan-Killiany ^98^. (**Extended Data Fig. 2**).

### Gene filtering by differential stability

Genes were filtered for those that showed more similar spatial patterns of expression across the six donors using the metric of differential stability (DS) as previously described by Hawrylycz *et al.* ^35^. For each gene, DS was calculated as the average correlation of that gene’s regional expression vector between each donor pair (15 pairs with all six brains, or 3 pairs in the triplets analysis, see below). Genes were ranked by DS and then only the top 50% percent of genes were retained. The 50% threshold was chosen on the basis of a grid-search (in combination with the region filter to optimise for generalisability) where the threshold for DS was varied between 10% and 100% (**Extended Data Fig. 1**).

### Filtering regions by donors represented

Regions were filtered for those that included samples from at least three of the six AHBA donor brains, which in the HCP-MMP1.0 parcellation retained 137/180 regions. Note that in the triplets analysis (see below), this means only brain regions with samples from all three donors in the triplet were retained. The choice to filter for representation of three of the six donors was chosen on the basis of a grid-search in combination with the differential stability gene filter to optimise for generalisability (**Extended Data Fig. 1**).

### Triplets analysis: disjoint triplet correlation as a proxy for generalisability

To test for generalizability we separated the six AHBA brains into pairs of disjoint triplets (for example donor brains 1,2,3 in one triplet and 4,5,6 in another). We applied our full analysis pipeline (including all processing steps e.g. probe aggregation, normalisation, and filters) independently to each of the twenty possible combinations of triplets, and correlated the regional scores for each DME or PCA component between each of the ten disjoint pairs (Pearson’s *r*). When filtering for consistently-sampled regions, the retained regions were different for each triplet of donor brains, so correlations were performed on only the intersection of regions retained in both triplets of each pair.

As the order of principal components can vary across different triplets, we employed a matching algorithm in which the full correlation matrix was computed between the top 5 principal components of both triplets (e.g. C1 from triplet A was correlated with each of C1-C5 of triplet B). The highest absolute correlation value in the matrix was then identified as representing two matched components and removed from the matrix, with the process repeated until all components were matched. The components were then ranked by the mean variance explained in each matched pair.

The median absolute correlation across all ten disjoint triplet pairs represented the generalisability, *g*, of the AHBA components processed using the given set of parameters. Processing parameters, in particular the filters for regions and donors, were optimised so as to maximise *g* while retaining as many genes and regions as possible; see **Extended Data Fig. 1**.

### Dimension reduction methods

Dimension reduction was performed using both principal component analysis (PCA) and diffusion map embedding (DME), the latter having been described for use in spatial gradient analysis of brain imaging data by Margulies *et al.* ^17^. For DME, the normalised cosine function was used as the kernel for the affinity matrix. No sparsity was added, and the alpha parameter was set at 1. These parameters were chosen as they optimised the inter-triplet correlation metric for generalisability. Both PCA and DME methods were implemented using the BrainSpace package ^99^. See Supplementary Methods for further explanation on DME and its benefits over PCA and other alternatives (e.g. ICA).

### Component gene weights

For each component, gene weights were computed as the Pearson correlation of each gene’s individual spatial expression vector with the regional scores of the component. For PCA these correlations are equivalent to the PCA loadings (eigenvectors) multiplied by the square root of the variance explained by the component (eigenvalues).

### Variance explained

For PCA, variance explained is given directly by the squared eigenvalues of the singular value decomposition. For DME, eigenvalues do not represent variance explained as the gene expression matrix is first converted to an affinity matrix using a kernel (here the normalized cosine). Therefore, variance explained was calculated as the difference in the total variance of the region-by-gene expression matrix before and after regressing the matrix on each component’s region scores.

That is, defining the residual regional expression vector of gene *g* after regressing out *i* components as *e_g,i_*, the total variance *V_i_* of the residualised region-by-gene expression matrix is

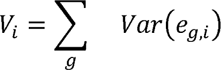

and for each component *C_i_*, variance explained *VE_i_* is given by

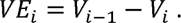

### Gene Ontology enrichment analysis for biological processes

Biological process enrichments of the gene weights for each component were computed using the ‘proteins with values/ranks’ function of online software *STRING* ^103^, which tests whether the mean weight of each annotated gene list is significantly higher or lower than random permutations of the same gene weights (the “aggregate fold change” method ^103,104^), and includes a Benjamini-Hochberg adjustment of the False Discovery Rate (FDR).

The aggregate fold change method was chosen as it does not require thresholding the gene weights of the components to define ‘target’ vs ‘background’ gene lists (as in e.g. Fisher’s exact test). That is, rather than setting a threshold for which genes are ‘in’ or ‘out’ of each component, we took the weighted gene list where all genes can have some contribution to each component, and for each component tested whether each Gene Ontology gene list was in aggregate more positively- or negatively-weighted than chance.

### Layer and cell-type enrichment analyses

The gene lists for cortical layer marker genes were obtained from published analyses of laminar enrichment in spatial transcriptomic data from human postmortem tissue in the dorsolateral prefrontal cortex ^38^ (columns Q-W of Table S4B in Maynard *et al.* ^38^).

Cell-type gene lists were obtained from Seidlitz *et al.* ^22^, who compiled lists of significantly differentially expressed genes from five independent single-cell RNA-seq studies ^44,105–108^. The gene list for synaptic marker genes was the unfiltered gene list from SynaptomeDB ^109^.

All enrichments for layers and cell-types were computed by the same aggregate fold change method ^104^ as in the *STRING* software ^103^, whereby the mean gene weight of each gene list was computed for both the true set of gene weights of each component, and for 5,000 random permutations of the weights. The *Z*-scores and permutation test *P*-values for significance testing of enrichment were corrected for multiple comparisons with the Benjamini-Hochberg FDR.

### GWAS enrichment analyses for educational attainment and intelligence

Genes associated with cognitive capacity by GWAS were obtained from:

- Lee *et al.* 2018, Supplementary Table 7 ^39^ (educational attainment).
- Davies *et al.* 2018, Supplementary Table 6 ^40^
- Savage *et al.* 2018, Supplementary Table 15 ^42^
- Hill *et al.* 2019, Supplementary Table 5 ^41^
- Hatoum *et al.* 2023, Supplementary Table 16 ^43^

Enrichment tests were performed by the aggregate fold change method ^104^, as above.

### Neuroimaging and other macro-scale brain maps (Fig. 2)

Neuroimaging and other macro-scale maps were obtained as follows:

- The 9 neuroimaging and macro-scale maps in the clustering analysis (**Fig. 2a**) were obtained from the *Neuromaps* package ^110^, and are also available in Sydnor *et al.* ^10^.
- The regions of cytoarchitectural differentiation (**Fig. 2b**) were obtained from Paquola *et al.* 2019 ^111^ and averaged into the HCP-MMP parcellation using the *Neuromaps* package ^110^.
- The map of fMRI degree (**Fig. 2c**) was obtained from Paquola *et al.* 2020 ^50^, and was originally computed from the HCP S900 release ^112^.
- The maps of MEG power bands (**Fig. 2d, Extended Data Fig. 5**) were obtained from the *Neuromaps* package ^110^.
- The map of adolescent change in cortical myelination was obtained from Váša *et al.* 2020 ^57^.

All maps were aggregated into HCP-MMP parcellation, and are provided in **Supplementary Table 3**.

Spatial associations between maps and the transcriptional components were computed by Pearson correlations and tested for significance using spin permutation tests (5,000 spins) by the Cornblath method ^113^, leveraging tools from *Neuromaps* ^110^, and tested for significance with FDR correction for multiple testing.

For the regions of cytoarchitectural differentiation, the mean component scores in each architectonic class were tested for differences between class mean scores using analysis of variance (ANOVA) against spin-permuted null models, followed by correction for FDR. The associations between individual cytoarchitectural regions and each component were computed by the Z-score of the mean component score in each region normalised by a spin permutation distribution of the regional mean component score with significance testing corrected for FDR.

### Single-cell co-variation analysis (Fig. 3a)

Single-cell RNA-seq data were obtained from the Allen Cell Types Database (https://portal.brain-map.org/atlases-and-data/rnaseq) ^58^.

Single-cell gene expression was filtered for the 7,873 genes in the optimally filtered AHBA dataset. To perform the analysis in **Fig. 3a**, the positive and negative gene weights were separated for each of C1-C3, and the dot product taken with the gene expression matrix of single-cell samples. This produced a vector of six numbers, representing the weighted total expression of C1+, C1-, C2+, C2-, C3+, C3-genes respectively, for each of the 50,000 single-cell samples.

That is, given the gene expression vector *s_j_* of each single-cell sample *j*, we computed the total weighted positive and negative expression *s*^+^*_j,C_i__*and *s*^−^*_j,C_i__* from the C1-C3 gene weights as:

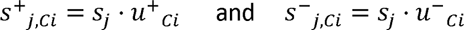

where *u^+^C_i_* = *max{u_Ci_, 0}* and *u^−^C_i_* = *min{uC_i_, 0}*.

### BrainSpan developmental gene expression processing (Fig. 3b-d)

BrainSpan data were obtained directly from the Allen Institute website ^6^ (http://brainspan.org) and processed as follows:

1. The 11 cortical regions in the BrainSpan data were manually matched to the HCP-MMP1.0 parcellation regions according to the descriptions in the BrainSpan documentation. This mapping is provided online at https://github.com/richardajdear/AHBA_gradients.
2. Exon-level expression data were filtered for only the matched BrainSpan regions.
3. Donor brains from which fewer than 4 regions were sampled were dropped.
4. Within each donor, expression of each gene was Z-normalised over regions.
5. Donors were aggregated into three age ranges (pre-birth, birth-13 years, and 18-40 years) and expression was averaged for each gene.

### AHBA-BrainSpan developmental consistency analysis (Fig. 3b-d)

Consistency between the AHBA components and BrainSpan was evaluated as follows:

1. Processed BrainSpan data were filtered for only the 7,973 genes retained in the filtered AHBA dataset (top 50% by differential stability; see above).
2. The dot product of the gene weights for C1-C3 were taken against the BrainSpan data, resulting in ‘BrainSpan scores’ for each of C1-C3, for each of the 11 BrainSpan regions, at each age range (pre-birth, birth-13 years, and 18-40 years).
3. In each of the 11 BrainSpan regions, ‘AHBA scores’ were computed as the mean of the matching HCP-MMP region scores from the original C1-C3 maps derived from the AHBA.
4. The ‘BrainSpan scores’ and ‘AHBA scores’ were correlated over the 11 BrainSpan regions (Pearson’s *r*), for each of C1-C3 and for each age bucket of the BrainSpan data.

As further clarification: given gene weights *u_i_* for AHBA component *C_i_* and the vector of expression over genes *b_j_* for each BrainSpan sample *j* (with a given age and region), the ‘BrainSpan score’ is

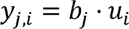

and the consistency was tested as the correlation across the matched regions of the AHBA scores *x* and the mean of the BrainSpan scores *y* of BrainSpan donors in each age range.

### BrainSpan developmental trajectory modelling (Fig. 3e)

The developmental trajectories of each decile of C1-C3 were computed as follows:

1. The ages in the BrainSpan data were converted to post-conception days on a log10 scale.
2. For each gene, a Generalised Additive Model was fitted using the *GLMGam* function in the *statsmodels* python package with alpha=1, and 12 3rd-degree basis splines as a smoothing function (df=12, degree=3 in the *BSplines* function). Sex and brain region were included as covariates.
3. Developmental curves were plotted from the fitted models for each gene, sex, and region, then averaged by decile of gene weight for each of C1-C3.

### Disorder spatial associations (Fig. 4a-b)

Maps of the regional centile score differences in cortical volume for ASD, MDD, and schizophrenia were obtained from the BrainCharts project by Bethlehem, Seidlitz, White *et al.* ^59^, in which normative models were computed for multiple brain phenotypes across the human lifespan from a harmonised dataset of >125,000 total MRI scans (N_controls_ = 38,839, N_ASD_ = 381, N_MDD_ = 3,861, N_SCZ_ = 315). As these data were in the Desikan-Killiany parcellation, the AHBA components in the HCP-MMP parcellation were mapped to a vertex-level surface map (FreeSurfer’s 41k fsaverage atlas) then re-averaged into the Desikan-Killiany parcellation. Pearson correlations with cortical maps of C1-C3 scores were computed, significance was assessed by spin permutation tests, and corrected for FDR across all nine tests (three disorders by three components).

These disorder maps are provided in **Supplementary Table 4**.

### Disorder DEG associations (Fig. 4c)

Differentially expressed genes (DEGs; FDR < 5%) from RNA-seq of postmortem brain tissue were obtained from the following case-control studies for each of ASD, MDD, and schizophrenia:

- ASD:

- Gandal *et al.* 2022, Supplementary Table S3 ^36^, WholeCortex_ASD_FDR < 0.05 l1 Gandal *et al.* 2018, Supplementary Table S1 ^65^, ASD.fdr < 0.05
- Parikshak *et al.* 2016, Supplementary Table S2 ^64^, FDR-adjusted P value, ASD vs CTL < 0.05
- MDD

- Jaffe *et al.* 2022, Supplementary Table S2 ^66^, Cortex_adjPVal_MDD < 0.05
- Schizophrenia

- Fromer *et al.* 2016, Supplementary Table S16 ^67^, FDR estimate < 0.05
- Gandal *et al.* 2018, Supplementary Table S1 ^65^, SCZ.fdr < 0.05
- Jaffe *et al.* 2018, Supplementary Table S9 ^70^, fdr_qsva < 0.05
- Collado-Torres *et al.* 2019, Supplementary Table S11 ^69^, adj.P.Val < 0.05 & region == ‘DLPFC’

A consensus list of DEGs was compiled for each disorder (except MDD where only one study was included) by including only those genes identified in at least 2 studies.

Enrichments for these gene sets in each disorder were computed by the aggregate fold change method ^104^, i.e. computing the percentile of the mean weight of the DEGs in C1-C3 relative to the 5,000 random permutations of the gene labels.

### Disorder-associated genes from GWAS (Fig. 4d)

Genes significantly associated with ASD, MDD, and schizophrenia by GWAS were obtained from:

- ASD: Matoba *et al.* 2020, Supplementary Table S7 ^60^
- MDD: Howard *et al.* 2019, Supplementary Table S9 ^61^
- Schizophrenia: Trubetskoy *et al.* 2022, Extended GWAS ^62^: https://figshare.com/articles/dataset/scz2022/19426775?file=35775617

Associations with GWAS were calculated using three methods (**Supplementary Figure S6**):

- Enrichment of the prioritised genes identified in each of the specific studies, using the aggregate fold change method ^104^ as described above.
- MAGMA ^71^, a regression technique which tests for association between each of the components C1-C3 and the *P*-values for each gene’s association with ASD, MDD or SCZ (from corresponding primary GWAS studies) without requiring a threshold to be applied to the GWAS-derived *P*-values to define a prioritised subset of genes for enrichment analysis. MAGMA additionally accounts for gene length and gene-gene correlations. The COVAR function of MAGMA was used to test for association of the GWAS *P*-values with the C1-C3 gene weights as a continuous variable. For standard MAGMA, a SNP-to-gene mapping window of +35kb/-10kb was used.
- H-MAGMA ^72^, an extension of MAGMA where SNP-to-gene mapping is performed using Hi-C chromatin measurements from postmortem brain tissue so as to capture trans-regulatory effects. We used the Hi-C mapping from adult brain DLPFC <u>available online</u> from the original H-MAGMA authors.

### Laminar enrichments shared across DEG and GWAS gene sets (Fig. 4f)

Enrichments for the marker genes of each cortical layer ^38^ were computed for the disorder-associated gene lists from DEGs and GWAS using Fisher’s exact test. These enrichments were computed both with and without filtering for only genes with positive C3 weights.

### Schizophrenia supragranular-specific cortical thinning (Fig. 4g)

The MRI-derived map of supragranular cortical thinning in schizophrenia was obtained from Wagstyl *et al.* ^63^ (N=90 subjects, 46 cases), and parcellated using the HCP-MMP1.0 parcellation. Pearson’s correlations were computed with C1-C3 and significance assessed by spin permutation tests, corrected for FDR.

## Data availability

Regional scores and gene weights for the transcriptional components C1-C3 are provided in **Supplementary Table 1**.

Gene expression datasets used are all publicly available:

- The Allen Human Brain Atlas is available at http://human.brain-map.org, and individual donor HCP-MMP parcellation images at https://figshare.com/articles/dataset/AHBAdata/6852911.
- The BrainSpan Atlas is available at https://www.brainspan.org/.
- The Allen Human Cell Atlas is available at https://portal.brain-map.org/atlases-and-data/rnaseq.
- The PsychENCODE dataset is available at https://github.com/dhglab/Broad-transcriptomic-dysregulation-across-the-cerebral-cortex-in-ASD.

Neuroimaging maps of healthy brain features are available in the neuromaps package (https://github.com/netneurolab/neuromaps). For convenience all brain maps used are provided in **Supplementary Table 3-4**. Gene lists used for enrichment analyses were all obtained from prior publications as detailed in **Methods**.

## Code availability

Analyses were performed with Python v3.10.5 and R v.2.2. Key python packages include: abagen==0.1.3, brainspace==0.1.10, neuromaps==0.0.3. Full details of all packages, a Dockerfile and link to docker image, and all code used for these analyses are publicly available at https://github.com/richardajdear/AHBA_gradients.

