## Supplementary Information for "Cortical gene expression architecture links healthy neurodevelopment to the imaging, transcriptomics, and genetics of autism and schizophrenia"

**Supplementary Methods**

1. Further details on dimension reduction

**Supplementary Tables**

1. C1-C3 region scores and gene weights *
2. GO terms table *
3. Normative brain maps *
4. Disorder brain maps *
5. Summary table of transcriptional datasets used for validation

* Provided in separate excel file

**Supplementary Figures**

1. Optimised AHBA components C1-C3 aligned with previously published PCs
2. Proportion of variance explained for C1-C3 was proportional to the number of strongly weighted genes
3. Enrichments for C1 aligned with prior results
4. Enrichments were not distinct for the less stable 50% of genes
5. Covariation in single-cells between positive and negative genes was only consistent for the same component, not between different components
6. GWAS associations were consistent when not thresholded for prioritised genes

### Supplementary Methods

#### Further details on dimension reduction

Diffusion map embedding (DME) is a graph Laplacian dimension reduction technique that can be understood as a form of non-linear PCA [100,101]. When applied to the region-by-gene expression matrix of the AHBA, PCA finds principal components by directly computing the eigenvectors of the matrix. In contrast, DME first projects the data into a region-by-region affinity matrix using a kernel (e.g. normalised cosine, Gaussian, correlation, etc), then finds successive components such that, in the low-dimensional space defined by the components, regions which are highly connected in the affinity matrix are close together and regions that are weakly connected are far apart.

In both PCA and DME, the regional scores of the components have zero correlation by construction. However, in PCA, components are also forced to have uncorrelated gene weights, i.e. the components have zero correlation between their eigenvector coefficients. DME relaxes this second constraint, resulting in components that are allowed to have some correlation in their genetic loadings. In practice this cross-correlation of gene weights is minimal ($r < 0.05$) but nevertheless allows for some flexibility that results in more robust components. There is also no prior evidence that transcriptional components should represent mutually uncorrelated sets of co-expressed genes, as mandated by PCA eigenvectors, so the less constrained weighting of genes in DME components may more accurately reflect the underlying transcriptional architecture. Importantly, PCA and DME components were in practice very highly correlated ($r > 0.9$ for each of PC1/C1, PC2/C2, PC3/C3; **Extended Data Fig. 1**), demonstrating that DME identifies broadly the same transcriptional components as PCA in a more robust and biologically plausible manner.

Independent component analysis (ICA) was also tested as an alternative to DME and PCA, but we found that ICA did not reproduce the well-known PC1 of the AHBA that has been extensively characterised in the prior literature [2,8,10]. ICA components were very different ($|r| < 0.4$) to the PCA and DME components, and also introduced additional uncertainty because: i) lower-order ICA components vary depending on the number of components to be estimated ; ii) ICA components do not have a natural ordering; and iii) the ICA algorithm uses a quadratic optimisation that depends on initial conditions [102]. In transcriptomics, ICA is most often used to isolate systematic sources of noise such as technical confounds, which is less applicable for our analysis with the AHBA where rows are not independent and identically-distributed samples but rather represent unique spatial locations on the cortex.

### Supplementary Tables

#### Table S5: Summary of transcriptional datasets used for validation

| **Dataset** | **N donors** | **# cortical locations** | **Analysis method** | **Validation result for C1, C2, C3 (*g* or *r*)** [a] |
|---|---|---|---|---|
| AHBA[1] (bulk microarray) | 6 | 2000+ (summarised to 180 regions) | DME, 50% most stable genes, regions with samples from 3+ donors | *g* = 0.97, 0.72, 0.65 |
| PsychENCODE[2] (bulk RNA-seq) | 54 | 11 | PCA, 50% most stable genes[b] | *r* = 0.85, 0.75, 0.73 |
| | | | PCA, all genes[c] | *r* = 0.83, 0.60, 0.56 |
| | | | DME, 50% most stable genes[d] | *r* = 0.82, 0.65, 0.64 |
| BrainSpan[3] (bulk RNA-seq) | 8[*] | 11 | Projection of AHBA C1-C3 gene weights[e] | *r* = 0.96, 0.88, 0.84 |
| Allen Cell Atlas[4] (single-cell RNA-seq) | 3 | 8 | Projection of AHBA C1-C3 gene weights: pos. vs neg.[f] | *r* = -0.90, -0.85, -0.95 |

*[a] For the AHBA, the validation was performed using the intra-dataset generalisability measure g, defined as the median regional correlation of components from disjoint triplets of donor brains.*
*[b] The 50% most stable genes were calculated by taking the healthy control subjects who had samples from at least 10 of the 11 cortical regions, then using the same differential stability method as for the AHBA, i.e. ranking genes by the mean correlation of spatial expression patterns across all pairs of subjects (12,418 genes for the PsychENCODE dataset). All RIN > 3 samples from N = 54 control subjects were then averaged into a {11 regions x 12,418 genes} matrix, to which PCA was applied.*
*[c] As in [b], but without filtering genes, i.e. PCA applied to the full matrix {11 regions x 24,836 genes}.*
*[d] As in [b], but using DME instead of PCA. DME first converts the {11 regions x 12,418 genes} matrix into a {11 regions x 11 regions} affinity matrix, so PCA is preferable for these low-granularity data.*
*[e] The Brainspan analysis examined whether C1-C3 were consistent at different donor ages. Therefore the gene weights from the AHBA-derived C1-C3 were projected onto the group-average Brainspan data, to examine if the AHBA gene profiles had consistent spatial expression patterns in Brainspan.*
*[f] The Allen Cell Atlas analysis examined whether C1-C3 were consistent within single cell nuclei. Therefore the gene weights from the AHBA-derived C1-C3 were projected onto each single-cell nuclei, separately for positively- and negatively-weighted genes. Here the validation results are (anti-)correlations between the positive and negative genes for each component across single cells (i.e. these correlations are over 50,000 cells, not correlations over the 8 sampled brain regions).*
*[*] 18-40 years subjects only; the same analysis was also performed for Pre-Birth and Birth-13y*

### Supplementary Figures

#### S1: Optimised AHBA components C1-C3 aligned with previously published PCs

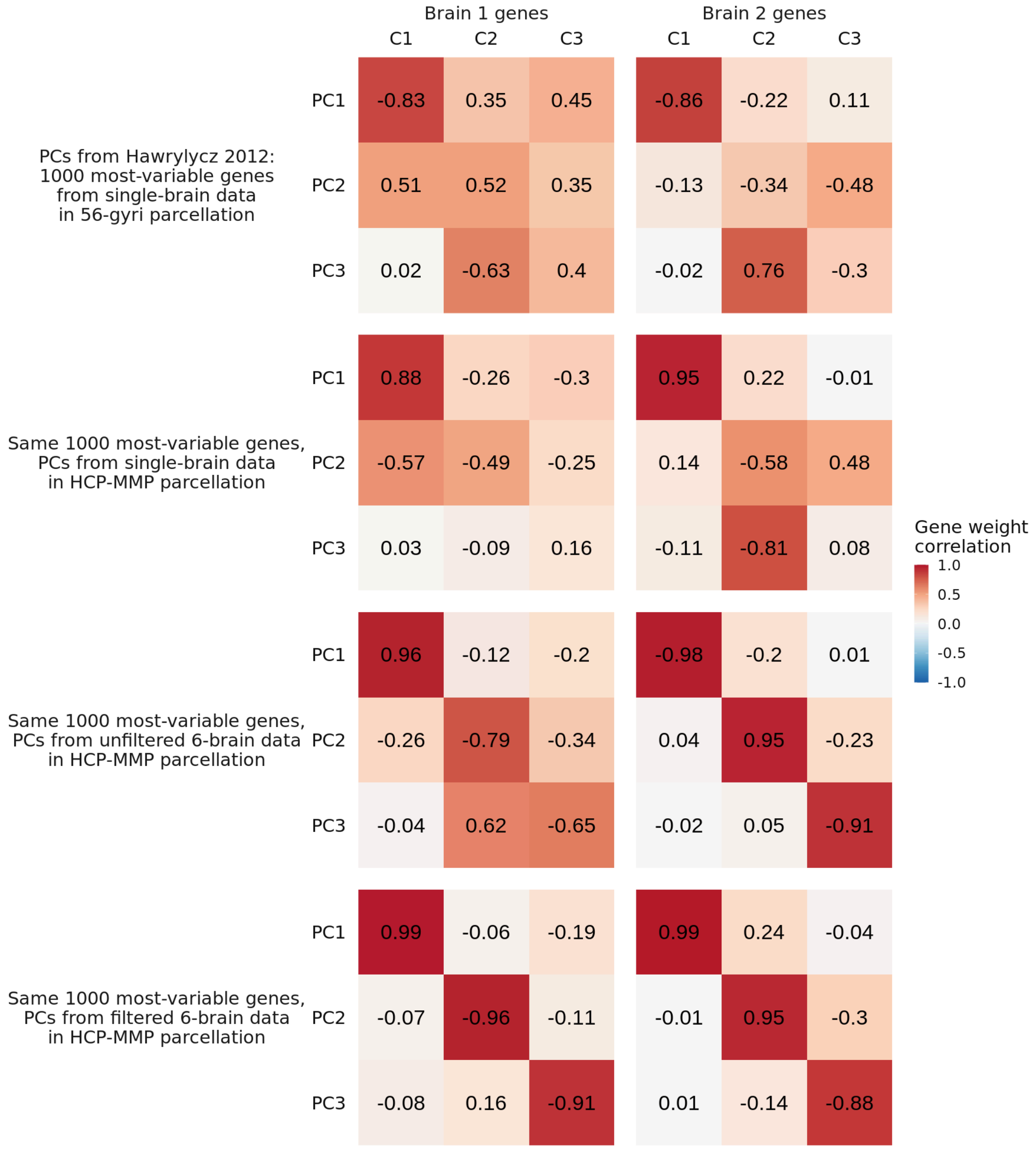


***Supplementary Figure S1: Optimised AHBA components C1-C3 aligned with previously published PCs.*** *AHBA components C1 and C2 are consistent with previously reported PCs computed from 1000 genes in single AHBA brains. In the initial release of the AHBA that included just two donor brains, Hawrylycz et al. showed that principal components of the expression of the 1000 genes with greatest spatial variation computed in a 56-region parcellation of a single brain were consistent between the two donor brains. Here we show that the gene weights of those PCs have similar alignment with the C1 and C2 that we derive using DME on 7973 genes across six donor brains in the HCP-MMP parcellation for different variations of data processing.*

#### S2: Proportion of variance explained for C1-C3 was proportional to the number of strongly weighted genes

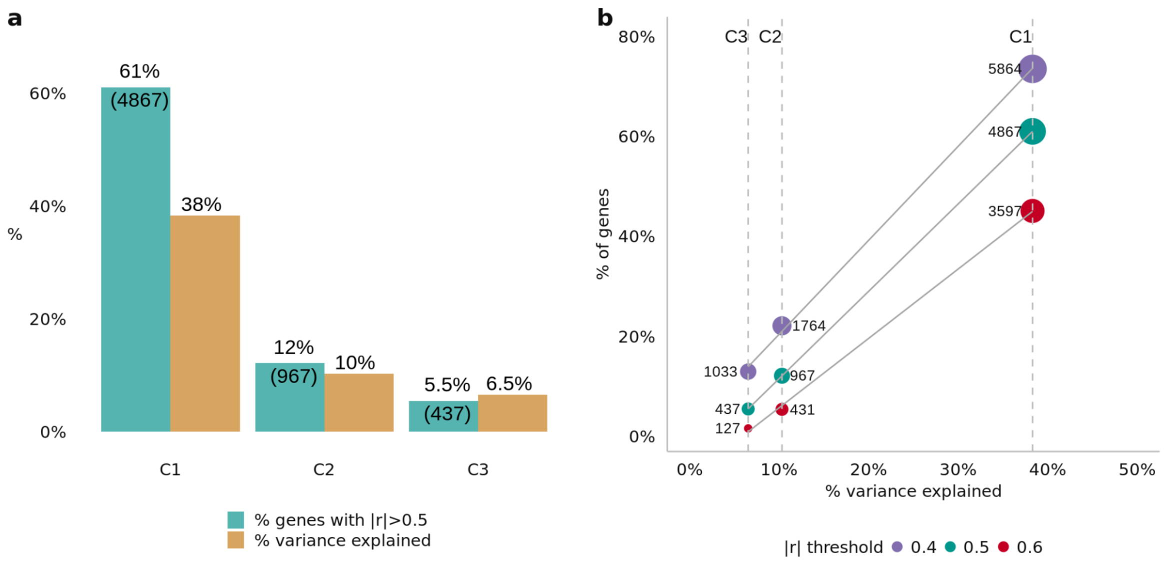


***Supplementary Figure S2: Proportion of variance explained for C1-C3 was proportional to the number of strongly weighted genes. a**, For each of C1-C3, the % of genes with absolute correlation |r|>0.5 is shown alongside the % variance explained (with the number of |r|>0.5 genes in parentheses), demonstrating that the % variance explained by higher-order components is related to the number of strongly weighted genes. **b**, The scatter plot shows the linear relationship between % variance explained and % strongly weighted genes for a range of threshold values (|r| > 0.4, 0.5, or 0.6). Size of points and their annotations display the number of genes above the given |r| threshold. For each |r| threshold, the slope of the linear relationship is >1, indicating that higher-order components C2 and C3 explain relatively more variance than expected given the number of strongly weighted genes alone (i.e. that C2 and C3 genes have relatively higher spatial variation in expression).*

#### S3: Enrichments for C1 aligned with prior results

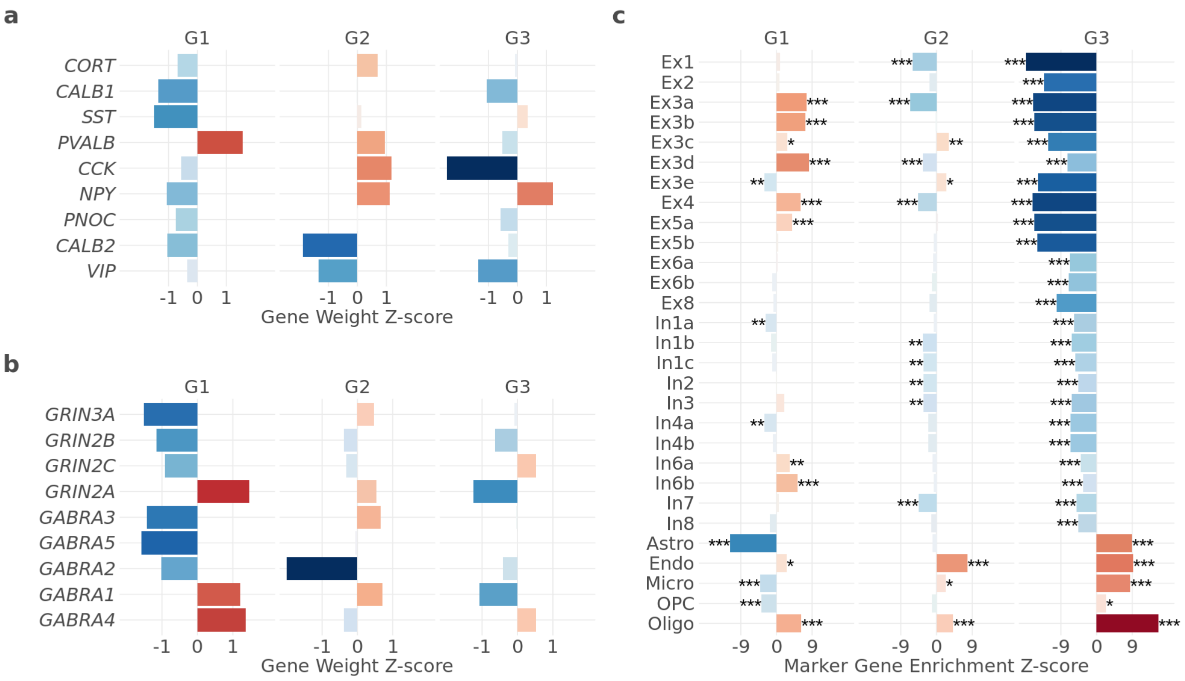


***Supplementary Figure S3: Enrichments for C1 aligned with prior results. a,*** *z-score of gene weight of key interneuron genes analysed for PC1 by Burt et al.* [5]. ***b****, as in* ***a*** *for key glutamatergic genes.* ***c****, enrichment (computed as z-score of mean gene weight of marker genes relative to permutations) of marker genes for neuron subclasses* [6]*. Significance was computed by two-sided permutation tests (Methods) and FDR-corrected across all tests; *, **, *** respectively indicate FDR-corrected two-sided p-value < 0.05, 0.01, 0.001.*

#### S4: Enrichments were not distinct for the less stable 50% of genes

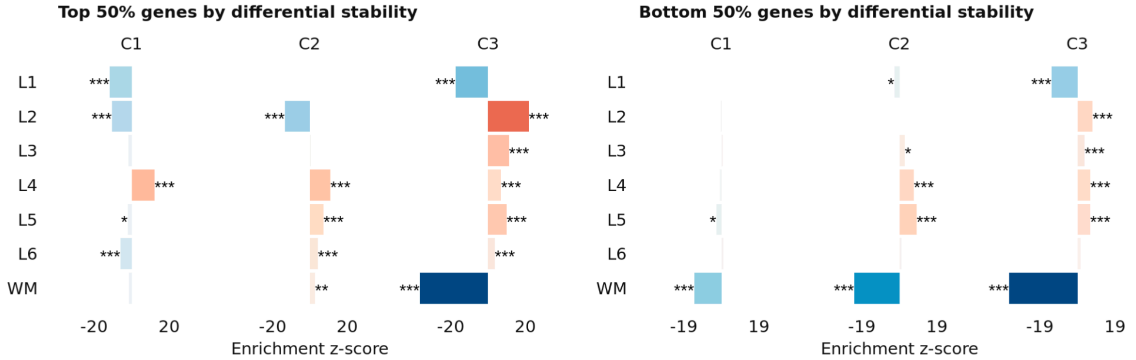


***Supplementary Figure S4: Enrichments were not distinct for the less stable 50% of genes.*** *Laminar enrichments from C1-C3 computed from the top 50% most stable genes (as in Fig. 1c-d), and the bottom 50% least stable genes. This demonstrates that the distinct laminar enrichments for C1-C3 depend on the gene stability filtering used to optimise the AHBA processing for generalisability. Significance was computed by two-sided permutation tests (Methods) and FDR-corrected across all tests; *, **, *** respectively indicate FDR-corrected two-sided p-value < 0.05, 0.01, 0.001.*

#### S5: Covariation in single-cells between positive and negative genes was only consistent for the same component, not between different components

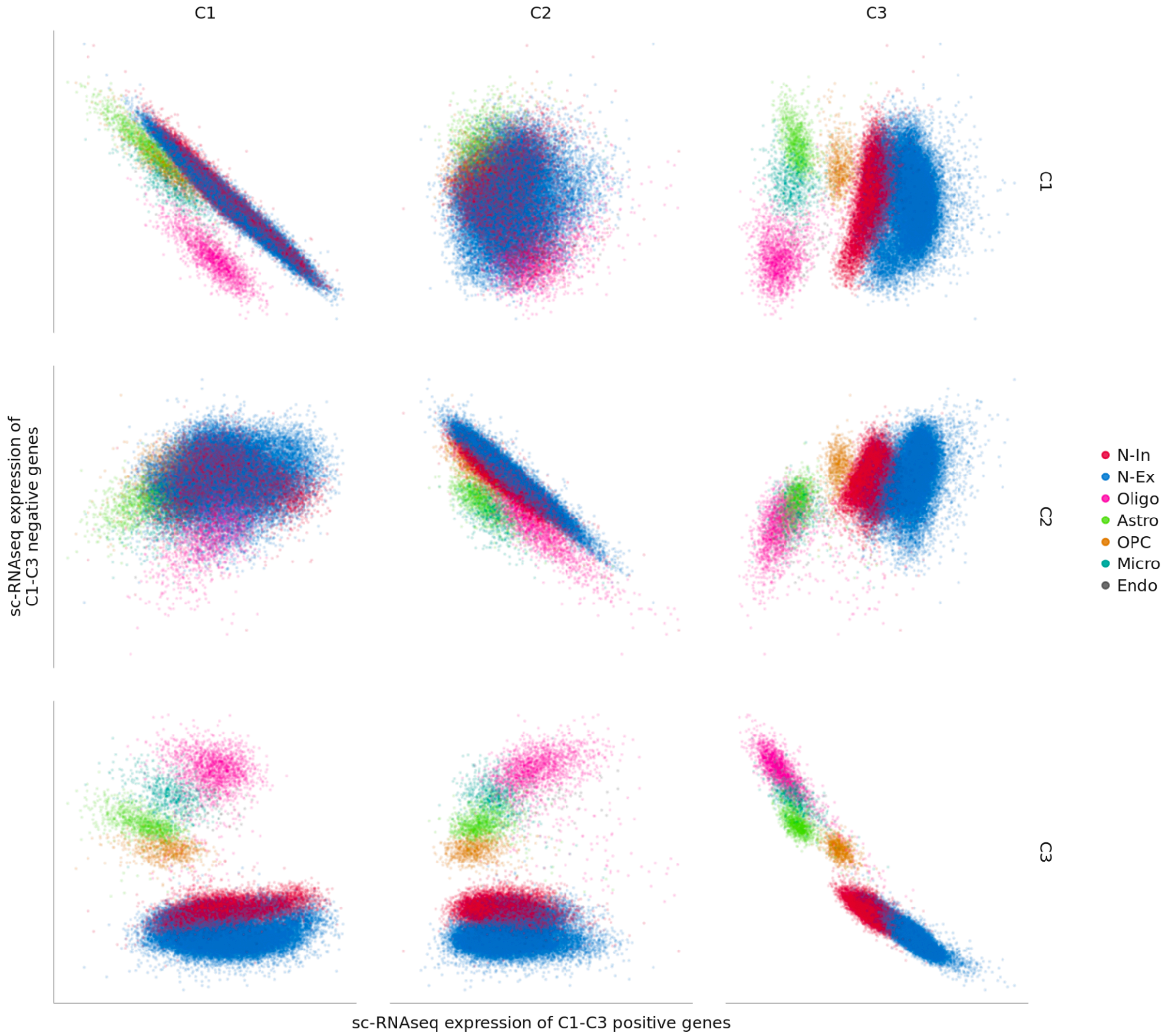


***Supplementary Figure S5: Covariation in single-cells between positive and negative genes was only consistent for the same component, not between different components.*** *This demonstrates that the observed anticorrelations are not a trivial result of e.g. total overall expression.*

#### S6: C1-C3 associations to autism and schizophrenia GWAS were consistent when not thresholded for prioritised genes

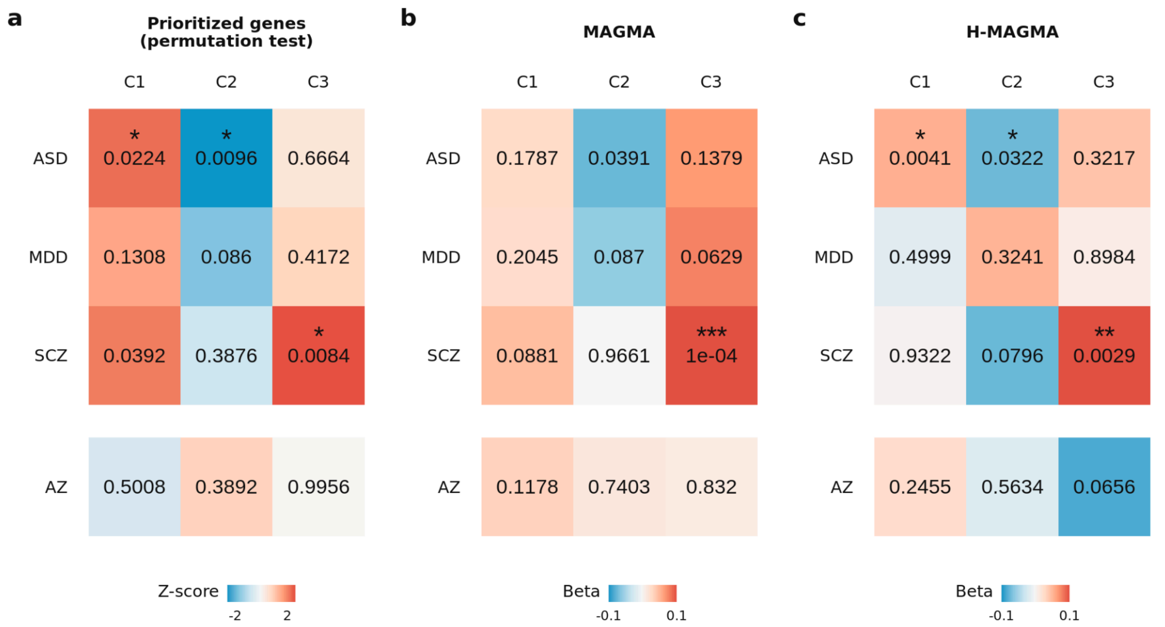


***Supplementary Figure S6: C1-C3 associations to autism and schizophrenia GWAS were consistent when not thresholded for prioritised genes. a**, Enrichment results for genes prioritised in the latest published GWAS studies of ASD [7], MDD [8], SCZ [9] (as in **Figure 4d**), and also Alzheimer's disease (AZ) [10], assessed using the aggregate fold change method [11,12]. Colour indicates the Z-score of the true mean C1-C3 weight of the prioritised genes relative to 5000 random permutations; numbers show the uncorrected two-sided p-values of this permutation test; *, **, *** indicate FDR-corrected two-sided p-value < 0.05, 0.01, 0.001, respectively. **b**, Associations for ASD, MDD, SCZ, and AZ from the same GWAS studies assessed using MAGMA [13], showing a strong association between SCZ and C3 consistent with the prioritised genes, but a weaker (though directionally consistent) association of ASD with C1 and C2. **c**, as in **b**, but assessed using H-MAGMA [14], which accounts for trans-regulatory genetic effects using Hi-C chromatin data from postmortem brain tissue. The use of H-MAGMA reproduced the significant association of ASD with C1 and C2, and the association of SCZ with C3. H-MAGMA was also the method used by Matoba et al. (2022) in the GWAS study to prioritise ASD-associated genes that were significantly associated with C1 and C2 (**a**). This suggests that trans-regulatory effects may play a significant role in how the spatially-expressed transcriptional programmes reflected in C1 and C2 are altered in ASD. Significance was computed by the MAGMA method [13] and FDR-corrected across all tests; *, **, *** respectively indicate FDR-corrected two-sided p-value < 0.05, 0.01, 0.001.*
